# Growth Factor-Based Manufacturing of Human Pluripotent Stem Cell-Derived Cardiomyocytes Using the Vertical Wheel Bioreactor System

**DOI:** 10.1101/2025.09.07.674777

**Authors:** Faisal J. Alibhai, Tamilla Valdman Sadikov, E. Coulter Montague, Lorena V. Cortes-Medina, Ian Fernandes, Gary Sun, Juliana Gomez-Garcia, Omar Mourad, Beiping Qiang, Sara S. Nunes, Gordon Keller, Michael A. Laflamme

**Author notes:** Corresponding Authors Faisal J. Alibhai, Michael A. Laflamme, University Health Network, 101 College Street, Rm 3-908, Toronto, ON Canada M5G 1L7.

## Abstract

**Background:** Multiple protocols have been reported for the large-scale generation of human pluripotent stem cell-derived cardiomyocytes (hPSC-CMs) in bioreactors using small molecules; however, no comparable bioreactor-based methods have been established using growth factors. This is despite evidence that differentiation with optimized concentrations of BMP4, Activin A, and bFGF offers finer control of cardiomyocyte phenotype. Here, we develop scalable hPSC expansion and growth factor-based cardiac differentiation protocols using the vertical wheel bioreactor (VWBR) system.

**Methods and Results:** The expansion of undifferentiated hPSCs was first optimized in 100 mL VWBRs by varying parameters, including starting cell seeding density, agitation rate, and media exchange schedule. Proliferation, viability, aggregate diameter, media metabolites, and pluripotency were assessed during hPSC expansion. Next, we evaluated the effects of undifferentiated hPSC culture conditions on subsequent cardiomyocyte differentiation potential. We found that hPSCs expanded in static culture or in VWBRs at different densities and agitation rates all differentiated into hPSC-CM populations of similar cardiac purity; however, cardiomyocyte yields were initially lower when VWBR-expanded hPSCs were used. We compared the differentiation kinetics of hPSCs expanded in VWBRs to conventional 2D culture and found that the former had accelerated mesodermal commitment and significantly greater cKit^+^/CXCR4^+^/PDGRFα^-^ cell formation during differentiation. Modifying our aggregation and mesoderm induction steps improved cell yields and enabled reliable production of >1x10^6^ cells/mL cardiac troponin T^+^ (cTnT) hPSC-CMs. Highlighting the versatility of our growth factor-based system, variation in the BMP4:Activin A ratio enabled a second heart field-like differentiation and generation of atrial-like cardiomyocytes in VWBRs. We further show that our expansion and differentiation protocols are reproducible and economical in 500 mL VWBRs, yielding on average 1.11x10^6^ hPSC-CMs/mL at a mean purity of 93% cTnT^+^. Characterization of VWBR produced hPSC-CM force generation, action potentials, and intercellular calcium transients confirmed the expected phenotype of ventricular-like cells. Lastly, VWBR produced hPSC-CMs robustly engrafted in the infarcted guinea pig myocardium, supporting use as a cell therapy product.

**Conclusions:** This novel bioreactor-based protocol will enable cardiac cell therapy and tissue engineering applications by providing scalable and consistent production of hPSC-derived cardiac cell products.

## Introduction

Ischemic heart disease is recognized as the leading cause of mortality worldwide. Up to 1 billion (1x10^9^) cardiomyocytes are lost during a myocardial infarction (MI; heart attack) involving a focal ischemic event in a region of the heart. Due to the adult heart’s poor regenerative capacity, the muscle lost following a MI is replaced by non-contractile scar tissue. Human pluripotent stem cells (hPSCs) are a promising source for the de novo generation of human cardiomyocytes (hPSC-derived cardiomyocytes; hPSC-CMs) that can be used to implant new heart muscle (i.e. remuscularize) within the infarcted myocardium. Small and large animal pre-clinical studies support this strategy, as transplanted hPSC-CMs form stable grafts that electromechanically couple with the host myocardium [1]. Large pre-clinical animal studies [2–4], as well as recent clinical trials [5], have involved the delivery of ≥100x10^6^ hPSC-CMs; therefore, scalable methods of hPSC-CM production are needed in order to meet these cell requirements.

Although hPSC culture and differentiation into cardiomyocytes has become straightforward at a small scale, further work is needed to establish efficient and economic expansion and differentiation protocols to supply sufficient cell quantities for downstream cell therapy and tissue engineering applications. Historically, scalable manufacturing platforms have relied on stirred tank bioreactors (STBs) to achieve larger-scale cell production. However, high shear stress domains within the vessel can potentially impact hPSC signalling and negatively affect production scale-up [5]. In contrast to established STB platforms, which use a horizontal impeller, the vertical wheel bioreactor (VWBR) is an alternative design that uses a vertical impeller to mix cells within the vessel. The advantage of this platform is the ability to achieve adequate cell mixing at substantially lower shear stress than can be achieved using STBs. This platform has been successfully used for the expansion of undifferentiated hPSCs using several commercially available lines [9–11], as well as the differentiation of cerebellar organoids and pancreatic beta-islet cells [6, 7].

With respect to hPSC-CM production, several protocols have been previously reported for the expansion and differentiation of hPSCs to hPSC-CMs using STB platforms [8–13]. These protocols have exclusively used small molecules to achieve mesoderm induction and subsequent cardiomyocyte specification. To date, no growth factor-based cardiomyocyte bioreactor differentiation protocol has been established, despite evidence supporting finer control over lineage specification compared to protocols that exclusively use small molecules [14, 15]. Here, we establish a scalable hPSC-CM production method using the VWBR as an all-in-one system for hPSC expansion and cardiomyocyte differentiation. By pairing expansion and differentiation using the same platform, we identified optimal hPSC expansion protocols for downstream cardiomyocyte differentiation. Identification of optimal conditions within the VWBR aided in the scaling of our differentiation protocol to the 500 mL VWBR scale and enabled the production of >500x10^6^ hPSC-CMs per vessel.

## Methods

### 2D static culture of undifferentiated hPSCs

Undifferentiated ESI-017 human embryonic stem cells (BioTime) were cultured on hESC-qualified Matrigel (Corning) coated tissue culture plastic in mTeSR1 medium (StemCell Technologies). Cells were passaged every 3-4 days using Accutase (EMD Millipore) and seeded into 10 cm dishes or T175 flasks at 18,000 cells/cm^2^ in mTeSR1 medium supplemented with the Rho-associated coiled-coil containing protein kinase (ROCK) inhibitor Y-27632 (10µM, StemCell Technologies). 24 hours after seeding, daily media exchanges were performed with mTeSR1 without Y-27632. hPSCs were confirmed to have a normal karyotype (Cytogenetics Laboratory, Hospital for Sick Children, Toronto, Ontario, Canada) and to be negative for mycoplasma using the e-Myco VALiD Mycoplasma PCR Detection kit (LiliF).

### 3D culture of undifferentiated hPSCs in VWBRs

100 mL and 500 mL VWBRs (PBS Biotech) were used for 3D hPSC expansion. After one passage in static 2D culture as described above, hPSCs were dissociated using Accutase (Thermofisher) and single cells were inoculated into VWBRs at the specified cell numbers in either mTeSR1 or StemScale (ThermoFisher) medium supplemented with 10 µM Y-27632. VWBRs were operated at full volume, 100 mL and 500 mL, with agitation rates as stated in revolutions per minute (rpm). For media exchanges, hPSC aggregates were allowed to gravity-settle for 5-8 minutes, after which the medium was aspirated from the vessel and replaced with fresh media. If aggregates did not settle in this time, the media was removed and centrifuged at 400 rpm for 3 minutes. Aggregates were resuspended in fresh medium without Y-27632 and returned to the VWBR vessel. To monitor cell density, aggregate diameter, and glucose/lactate levels, a 1 mL sample was removed from the VWBR daily, immediately prior to media exchanges when exchanges were performed. Aggregates were dissociated using Accutase, after which total cell number and viability (Acridine orange/propidium iodide, Revvity) were measured using the Cellometer K2 automated cell counter. The aggregate diameter for each day was measured manually using Image J, with a minimum of 80 aggregates for each replicate. On the final day of undifferentiated hPSC expansion, cells from 100 mL and 500 mL VWBRs were collected into a 50 mL or 250 mL conical tube, pelleted at 400 rpm for 3 minutes and resuspended in 10 mL Accutase supplemented with DNase I (∼20 U/mL; Millipore-Sigma) for 10 minutes at 37°C, pipetting the mixture once at the halfway point of the dissociation. Following enzymatic dispersion, cells were washed with 15 mL of DMEM/F12 media supplemented with DNase I and centrifuged at 1200 rpm for 5 minutes. Cell pellets were resuspended in DMEM/F12 medium, filtered through a 70 µm filter, and viable cells were quantified using the Cellometer K2 cell counter. Fold expansion was calculated by dividing the cell count at harvest by the starting cell density. Media glucose and lactate levels were measured by the Laboratory Medicine Program at Toronto General Hospital (Toronto, Ontario, Canada).

### Generation of ventricular cardiomyocytes

hPSCs were differentiated into ventricular CMs using a modified version of a previously reported growth factor-based protocol [16, 17]. Following dissociation of either 2D static or 3D cultured hPSCs, cells were washed, filtered through a 70µm filter, counted using an automated cell counter, and transferred to aggregation media consisting of StemPro-34 media (ThermoFisher Scientific) supplemented with L-glutamine (2 mM, ThermoFisher Scientific), L-ascorbic acid (50 μg/mL, Sigma), monothioglycerol (0.004% v/v, Sigma), and transferrin (150 μg/mL, Roche), hereafter referred to as 100% StemPro. On day 0, cells were aggregated in 100% StemPro medium supplemented with bone morphogenetic protein-4 (BMP4; 1 ng/mL, R&D Systems) and either Y-27632 (10 μM) or Chroman-1 (50 nM; Tocris Bioscience). For all conditions, cells were aggregated at 1x10^6^ cells/mL. Cells were aggregated at 70 rpm in poly-HEMA-coated 6-well plates on an orbital shaker or at a range of speeds in the VWBR, as indicated for each figure. 18 hours following aggregation, mesoderm was induced using 100% StemPro supplemented with BMP4, Activin A (R&D Systems), and basic fibroblast growth factor (bFGF; 5 ng/mL, Peprotech). BMP4 and activin A concentrations were varied as described in their respective figures. For a subset of experiments, the glycogen synthase kinase (GSK) 3 inhibitor/Wnt activator CHIR99021 (1 µM, StemCell Technologies) was also added during the mesoderm induction stage from day 1-3. Unless otherwise stated, mesoderm induction was allowed to occur for 48 hours. On day 3, aggregates were washed once with Iscove’s Modified Dulbecco’s Medium (IMDM) media and resuspended in 100% StemPro supplemented with the Wnt inhibitor IWP2 (2 μM, Tocris Bioscience) and vascular endothelial growth factor (VEGF; 10 ng/mL, Peprotech) and cultured for an additional 3 days. On day 6, an 80% media exchange was performed using medium consisting of 25% (v/v) StemPro-34 and 75% (v/v) IMDM, supplemented with 2 mM L-glutamine, 50 μg/mL L-ascorbic acid, 0.004% monothioglycerol, and 75 μg/mL transferrin; hereafter referred to as SP25 medium. During culture on days 6-10, SP25 medium was also supplemented with 5 ng/mL VEGF and changed 80% every other day. After day 10, aggregates were maintained in SP25 medium without VEGF, with 80% media exchanges every other day. For a subset of experiments, aggregates were transferred on day 12 into RPMI-1640 (ThermoFisher Scientific) medium supplemented with B27 (ThermoFisher Scientific) and 2 mM L-glutamine, followed by media exchanges every other day. All guided differentiation protocols were performed in a standard cell culture incubator (5% CO_2_ and 21% O_2_) in suspension using Poly-HEMA (Sigma) coated 6-well plates, 100 mL VWBRs, or 500 mL VWBRs.

### Generation of atrial cardiomyocytes

To generate atrial-like cardiomyocytes, we used a modified version of the preceding differentiation protocol [18]. For this, hPSCs were aggregated as previously described. On day 1, cultures were treated with a mesoderm induction media consisting of 100% StemPro supplemented with 4 ng/mL BMP4, 2 ng/mL Activin A, and 5 ng/mL bFGF. On day 3, cells were washed with IMDM and cultured in 100% StemPro supplemented with 2 µM IWP2, 10 ng/mL VEGF, and 500 nM all-trans-retinoic acid (Sigma). On day 5, cultures were transferred into SP25 medium supplemented with 5 ng/mL VEGF. After day 10, cultures were maintained in SP25 medium without VEGF with 80% media exchanges every other day.

### hPSC-CM dissociation and cryopreservation

hPSC-CM aggregates were harvested at the indicated time points and enzymatically dispersed using 300 U/mL collagenase II (Worthington) in Hanks Balanced Salt Solution (Wisent) overnight at room temperature. The following day, cells were washed and filtered through a reversible cell strainer (Stem Cell Technologies). Any remaining aggregates were further dissociated using TrypLE (ThermoFisher) supplemented with 20 U/mL DNase I for 3 minutes at 37°C. Cells were washed with SP25 medium supplemented with DNase I, then resuspended in SP25. For cryopreservation, cells were resuspended in CryoStore CS10 (Biolife Solutions) and frozen using a controlled rate freezer (BioCool) operated at a cooling rate of 1°C/min from 0°C to -50°C. Cells were then transferred to a -80°C freezer overnight, after which cells were transferred to a -150°C freezer for long-term storage.

### Flow cytometry

To assess undifferentiated hPSC pluripotency on the final day of expansion in the VWBR, aggregates were dissociated using Accutase, single cells fixed and permeabilized using the Cytofix/Cytoperm kit (BD Biosciences), and cells stained with primary conjugated antibodies against Oct4 and Sox2. To assess mesoderm induction, aggregates were collected on day 2, 2.5, 3 or day 4 of differentiation, dispersed to single cells using TrypLE, washed with FACS buffer, and stained with primary antibodies for 30 minutes at room temperature. For atrial-like cardiomyocyte differentiations, day 4 cells were stained for CD235a and aldehyde dehydrogenase activity using the ALDEFLUOR™ kit (StemCell Technologies) according to the manufacturer’s instructions. To assess hPSC-CM purity, hPSC-CMs were fixed with 4% (w/v) paraformaldehyde (PFA) for 10 minutes, permeabilized with ice-cold 90% methanol for 20 minutes, and stained with primary antibodies against cardiac troponin T (cTnT, Miltenyi Biotec), myosin light chain-2v (MLC2v, Miltenyi Biotec), and myosin light chain-2a (MLC2a, Miltenyi Biotec) for 30 minutes at room temperature. Events were acquired using an LSRII or Fortessa flow cytometer. All antibodies and concentrations used are listed in **Supplementary Table 1**. Data were analyzed using FlowJo (Tree Star).

### Immunofluorescence staining

hPSC or hPSC-CM aggregates were harvested and fixed in 4% PFA for 30 minutes at room temperature. Following two PBS washes, aggregates were embedded in 2% agarose or HistoGel™ (ThermoFisher), then submitted for standard histological processing, paraffin embedding, and sectioning at 5 µm intervals. Sections were deparaffinized and rehydrated using a series of xylene and alcohol washes. Heat-induced epitope retrieval was performed using 10 mM sodium citrate (pH 6.0) with 0.05% Tween 20. Aggregate sections were blocked with 10% normal goat serum for 1 hr at room temperature, followed by immunostaining with unconjugated primary antibodies overnight at 4°C. All primary antibodies are listed in **Supplementary Table 1**. Sections were washed and stained with secondary antibodies for 1 hr at room temperature. The following secondary antibodies were used: goat anti-mouse IgG Alexa Fluor488 (ThermoFisher), goat anti-rabbit Alexa Fluor594 (ThermoFisher), and goat anti-rabbit Alexa Fluor647 (ThermoFisher). Following secondary antibody staining, DAPI (Biotium) was applied for 10 min at room temperature. Sections were washed and then mounted using ProLong™ Diamond Antifade Mountant (ThermoFisher). Histological sections of engrafted hearts were prepared as described previously [17]. In brief, 14 days after cell transplantation, hearts were harvested and fixed with 10% neutral buffered formalin for 48 hours, after which hearts were transversely sectioned at 3 mm intervals using a heart matrix (Zivic), and paraffin-embedded for immunohistochemical analyses. Sections (4 µm thick) underwent heat-induced antigen retrieval in citrate buffer and were immunostained with anti-α-actinin and anti-Ku80. Ku80 was detected using a biotinylated goat anti-rabbit secondary antibody (Vector Labs) for 1 hour at room temperature, followed by detection using the ABC-AP kit (Vector Labs) and Vector Red (Vector Labs) according to the manufacturer’s instructions. For visualization of α-actinin, sections were incubated with an Alexa-647 fluorophore-conjugated secondary antibody for 1 hr at room temperature. Sections were counterstained with a FITC-conjugated collagen hybridizing peptide (F-CHP, 3Helix) to label scar tissue. Sections were scanned using a Zeiss AxioScan fluorescence scanner with a 20x objective (Plan Apochromat 0.80 NA) by the Advanced Optical Microscopy Facility (University Health Network, Toronto), and the graft region was quantified using QuPath.

### Gene expression analysis

Total RNA was isolated using the RNAqueous-micro kit (Invitrogen) with post-column DNase treatment or the miRNeasy RNA isolation kit (Qiagen). cDNA synthesis (iSCRIPT, BioRad) was performed using up to 1 µg of RNA. Subsequent RT-qPCR was performed using a CFX Real-Time instrument (BioRad) and either the SensiFAST SYBR No-ROX SYBR Green (Bioline), QuantiFast SYBR Green PCR kit (Qiagen) or Advanced Universal SYBR Green Supermix (BioRad), with primers as described in **Supplementary Table 2**. Experiments comparing ventricular and atrial-like cardiomyocyte gene expression are presented relative to TBP. Gene expression was measured using technical duplicates, evaluated for relative copy number, reaction efficiency, and genomic DNA contamination (<0.01% of TBP content), using a 10-fold dilution series of sonicated human genomic DNA standards made in-house from wild-type HES2 hESC cells, ranging from 2.5 pg/mL to 25 ng/mL, as described previously [19]. Gene expression was otherwise quantified by the 2^-ΔΔCT^ method using the mean expression of TBP and DDB1 as housekeeping genes.

### Optical Mapping

Dual voltage and calcium optical mapping were performed as described previously [20]. Briefly, following collagenase dissociation of day 20 hPSC-CM aggregates, single cells were plated onto Matrigel coated 6 well dishes at 2x10^6^ cells per well in RPMI+B27 medium supplemented with 10 µM Y-27632. The following day, medium was replaced with fresh RPMI+B27 and exchanged every other day. One week later, hPSC-CMs were dissociated using 0.125% trypsin-EDTA (Corning) supplemented with 20 U/mL DNase I followed by inactivation with defined trypsin inhibitor (ThermoFisher). 1x10^5^ cells were plated onto Matrigel-coated 18 mm round glass coverslips (Fisher Scientific) in RPMI+B27 media supplemented with 10 µM Y-27632. The following day, medium was replaced with fresh RPMI+B27 and monolayers were allowed to form for an additional 2 days. 3 days after plating, monolayers were stained with 1 µM BeRST1 [21], and 2.5µM Calbryte^TM^ 520 AM in RPMI+B27 media for 1 hour at 37°C in the dark. Optical action potentials and intercellular calcium transients were acquired and analyzed using the protocol described previously [20]. Signals were acquired during spontaneous beating and following point electrical stimulation at 1, 1.5, and 2 Hz.

### Contractile measurements using biowire preparation

Biowire cardiac tissues were prepared using hPSC derivatives as previously reported [22, 23]. In brief, hPSC-CMs at differentiation day 20 were combined with human cardiac fibroblasts (Lonza) at a 3:1 ratio, and a total of 214,000 cells were seeded per device. Tissue constructs were cultured for 48 hours in RPMI+B27 supplemented with 10 µM Y-27632 and 1X antibiotic-antimycotic (ThermoFisher), after which tissues were transferred to 100% StemPro-34 medium supplemented with 1X antibiotic-antimycotic, and medium exchanges were performed every 2-3 days. On day 15, cardiac tissues were placed into a custom electrical stimulation chamber consisting of two parallel 3 mm diameter carbon rods (Ladd Research) spaced 1.5 cm apart. Stimulation chambers were placed under a microscope, and the platinum wire was connected to an Astro-Med Grass S88X stimulator. Measurements of active force of contraction normalized to cross-sectional area were performed as described previously [22, 23].

### Cardiac injury and hPSC-CM transplantation

All animal studies were reviewed and approved by the Animal Care Committee of University Health Network. To induce cryoinjury, 650-750 g male Hartley guinea pigs were anesthetized with ketamine (IP, 35 mg/kg) and xylazine (IP, 2 mg/kg), intubated, mechanically ventilated (60 breaths per minute, tidal volume of 5 mL/kg) and maintained with 1.5 % isoflurane in oxygen. At the time of anesthesia animals were also injected with slow-release buprenorphine (SQ, 0.3 mg/kg). A left thoracotomy was performed, the heart exposed, and cardiac cryoinjury induced using a 7.5 mm diameter, liquid nitrogen-cooled aluminum cryoprobe to the anterolateral LV four times for 30 seconds each. During recovery after the procedure, animals were given meloxicam (SQ, 0.3 mg/kg), Baytril (SQ, 5 mg/kg), and saline (SQ, 5 cc). At 10 days post-injury, a repeat thoracotomy was performed, and the injury site was directly injected with 70x10^6^ hPSC-CMs suspended in a previously reported pro-survival cocktail [24]. The pro-survival cocktail consisted of growth factor-reduced Matrigel supplemented with 200 nM Cyclosporine A (Sandimmune/Novartis), 50 µM Pinacidil (Sigma), and 100 ng/ml IGF-1 (Peprotech). To prevent xenograft rejection, all animals were treated with cyclosporine A (SQ, 15 mg/kg/day for 7days, followed by 7.5 mg/kg/day thereafter) and methylprednisolone (IP, 2 mg/kg/day), starting 2 days prior to cell transplantation and continuing until euthanasia. ***Statistical analyses***

All figures are presented as mean ± standard error of the mean (SEM). Linear regressions are displayed with 95% confidence intervals. Two-sample comparisons were made using a two-tailed unpaired Student’s t-test. Where stated, comparisons between multiple groups were analyzed using one- or two-way ANOVA, with multiple comparisons post-hoc tests indicated. All data were tested for normal distribution and equal variance. GraphPad Prism software (Version 10.4.1) was used for all statistical analyses, with the threshold for significance set at p≤0.05.

## Results

### Optimization of undifferentiated hPSC expansion in VWBRs

We first set out to establish a protocol for the expansion of undifferentiated hPSCs in VWBRs that would be compatible with downstream cardiomyocyte differentiation. Toward this goal, we assessed the impact that the initial cell seeding density, bioreactor agitation rate, and media exchange schedule had on hPSC expansion in the VWBR, with the goal of achieving at least 1x10^6^ viable cells/mL at the end of the expansion period. Based on prior work by other investigators [5, 25], we first tested different seeding densities in 100 mL VWBRs, at 2x10^4^, 8x10^4^, or 15x10^4^ cells/mL, while fixing the expansion period to 6 days and culture agitation rate to 40rpm (**Figure 1A**). For all three cell seeding densities, the hPSCs aggregated efficiently and expanded over time (**Figure 1B**), although cultures starting at lower cell densities resulted in lower cell yields on day 6, as anticipated. Cell viability was comparatively low on day 1, particularly for the 2x10^4^ cells/ml condition, but viability improved over time and was >90% for all three conditions at harvest on day 6 (**Figure 1C**). Importantly, hPSCs expanded in VWBRs under all conditions maintained the expression of the pluripotency markers Oct4, Sox2, Tra-1-60, and Tra-1-81 (**Supplementary Figure 1A-B**). hPSC cultures seeded with the lowest initial cell density (2x10^4^/mL) exhibited the largest fold expansion over 6 days (**Figure 1D**), but the final cell yield with this condition was lower than our target cell number. In contrast, VWBRs seeded with either 8x10^4^ or 15x10^4^ cells/mL yielded >100x10^6^ total cells at day 6. However, reactors seeded with 15x10^4^ cells/mL exhibited a plateau in glucose consumption (**Figure 1E**) and lactate production (**Figure 1F**) as early as day 4, implying saturation of the culture vessel. Moreover, the mean aggregate diameter was significantly larger with the 15x10^4^ cells/mL condition (**Figure 1G**), with maximum aggregate diameter exceeding 600 µm by day 4 of the expansion period (**Figure 1H-I**). hPSC aggregate diameter at harvest on day 6 was not significantly different for cultures initially seeded at 2x10^4^ and 8x10^4^ cells/mL, suggesting a threshold for which inputting more starting hPSCs can increase the number of cell aggregates formed without substantially affecting final aggregate diameter (**Figure 1G-I**). Based on these results, we decided to employ a starting cell density of 8x10^4^ cells/mL for subsequent experiments, as this condition showed a good balance between fold-expansion, yield, and final aggregate diameter.

**Figure 1.**
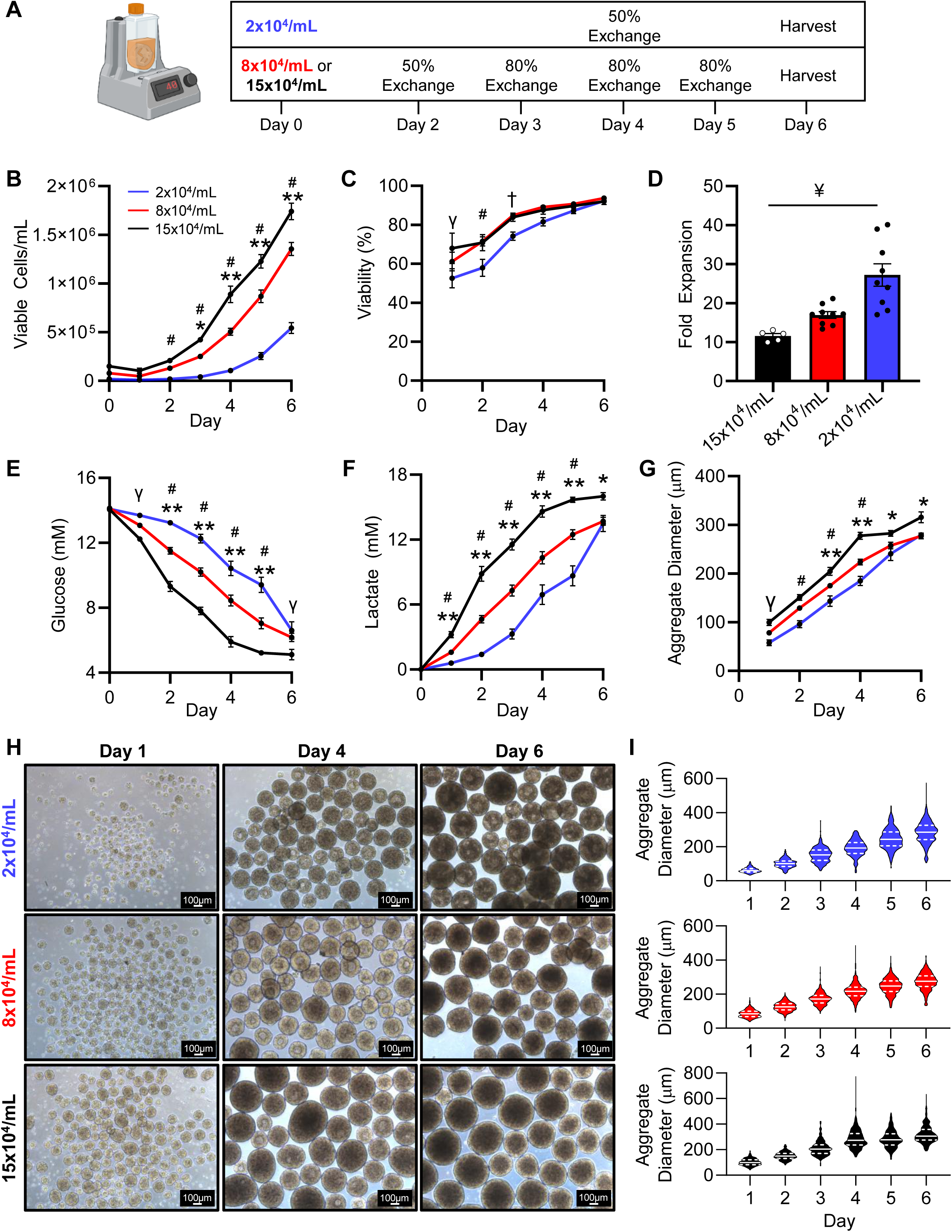
Inoculation density impacts hPSC expansion and yields in the 100mL VWBR. A) Schematic illustrating media exchange schedules for hPSC expansion following seeding with three cell densities (2x10^4^, 8x10^4^, or 15x10^4^ cells/ml). Cells were cultured at 40 rpm in mTeSR1 for 6 days. Cell density within the VWBR by condition as measured by the number of viable cells per mL (B) and % viable cells (C). D) Fold expansion of hPSCs over 6 days in the VWBR. E) Media glucose and F) lactate levels during the 6-day expansion protocol. G) Mean aggregate diameter, H) representative aggregate images, and I) aggregate diameter distribution during expansion in the VWBR. ^#^p≤0.05 2x10^4^ cells/mL vs. all other groups, *p≤0.05 15x10^4^ cells/mL vs. all other groups, **p≤0.01 15x10^4^ cells/mL vs. all other groups, ^†^p≤0.05 8x10^4^ cells/mL vs. 2x10^4^ cells/mL by two-way ANOVA followed by a Tukey post hoc. ^¥^p≤0.05 2x10^4^ cells/mL vs. all groups by one-way ANOVA followed by a Tukey post hoc. ^γ^p≤0.05 2x10^4^ cells/mL vs. 15x10^4^ cells/mL by two-way ANOVA followed by a Tukey post hoc. N=5-9/group.

Next, we tested a range of bioreactor agitation rates to determine how this parameter affects undifferentiated hPSC expansion. For this, 100 mL VWBRs were initially seeded with hPSCs at 8x10^4^ cells/mL, which were then cultured at either 30, 40, or 60 rpm (**Supplementary Figure 2A**). hPSCs maintained at lower agitation rates exhibited more rapid expansion (**Supplementary Figure 2B**); however, due to aggregate pooling at the bottom of the vessel, we had to harvest cultures maintained at 30 rpm on day 5, while cultures at 40 and 60 rpm could be harvested on day 6. Higher agitation rates were associated with poorer aggregation and lower cell viability on day 1, especially for cultures at 60 rpm. However, at the end of the expansion period all three conditions exhibited >90% cell viability and final cell densities of ≥1x10^6^ cells/mL (>13-fold expansion) (**Supplementary Figure 2B-D**). All three culture conditions showed similar glucose consumption and lactate production (**Supplementary Figures 2E-F**), although these parameters showed modestly reduced saturation for cultures maintained at 60 rpm, consistent with the slightly lower hPSC yield with this condition. As anticipated, the most striking effect of agitation rate was on hPSC aggregate diameter, with the 60 rpm cultures showing significantly smaller aggregates throughout the expansion process (**Supplementary Figure 2G-I**).

Next, we examined the effect of media exchange schedule on hPSC expansion. For this, we compared the media exchange schedule that had been employed in the preceding experiments (80% exchange after day 3) to two other protocols in which either 20% or 50% of the media was exchanged after day 3 (**Supplementary Figure 3A**). Interestingly, hPSC growth kinetics were similar for the 50% and 80% media exchange protocols, but cell yields were significantly lower for cultures undergoing 20% media exchanges (**Supplementary Figure 3B-D**). While it did not reach statistical significance, there was a trend toward higher viability at later time-points in cultures with 80% media exchanges (**Supplementary Figure 3C**). Moreover, cultures undergoing 50% media exchanges showed significantly lower glucose and higher lactate levels than the 80% media exchange condition at later time-points, reflecting reduced media replenishment despite similar cell densities (**Supplementary Figure 3E-F**). Despite the differences in hPSC expansion, all three protocols yielded hPSC aggregates with similar diameters, and all maintained pluripotency marker expression (**Supplementary Figure 3G-I**). Based on these data, we employed the protocol with 80% daily media exchanges after day 3 for all subsequent experiments, as this condition maximized hPSC viability and yield, while minimizing lactate accumulation.

While the previous experiments all involved hPSCs cultured in mTeSR1 medium, we also tested another commercially available medium, StemScale, for compatibility with hPSC expansion in the VWBR system. For this, we seeded VWBRs with hPSCs at 8x10^4^ cells/mL in either StemScale or mTeSR1, followed by culture at 40 rpm with media exchanges as depicted in **Supplementary Figure 4A**. Interestingly, hPSCs cultured in StemScale expanded more rapidly, reaching the target cell density by day 4 (**Supplementary Figure 4B-D**). Although cell viability on day 1 was higher for hPSC cultures in StemScale, likely due to better initial aggregation efficiency, cell viability was similar at the end of the expansion protocol for both media (**Supplementary Figure 4C**). Glucose levels were significantly higher in StemScale versus mTeSR1 cultures at early time-points, but this later equilibrated (**Supplementary Figure 4E**). Moreover, StemScale cultures had significantly higher lactate levels throughout, consistent with more rapid expansion than in mTeSR1 cultures (**Supplementary Figure 4F**). Lastly, hPSC aggregates were significantly larger in StemScale than in mTeSR1 and exhibited greater variability on the day of harvest compared to aggregates obtained from mTeSR1 cultures (**Supplementary Figure 4G-I**). Taken collectively, these data demonstrate that undifferentiated hPSCs can be efficiently expanded in VWBRs by optimizing parameters including initial seeding density, agitation rate, and medium selection.

### Testing the cardiac potential of undifferentiated hPSCs after expansion in VWBRs

After establishing reliable protocols for the expansion of undifferentiated hPSCs in VWBRs, we next examined whether hPSCs generated in this system would be compatible with our previously reported guided cardiac differentiation protocol [16, 17]. To test this, undifferentiated hPSCs were expanded in mTeSR1 medium in static 2D culture or VWBRs across a range of cell densities and agitation rates. The resultant hPSCs were then dispersed into single cells and subjected to our guided differentiation protocol in 6-well plates (**Figure 2A**). PDGFRα and CD56 were used as markers of cardiac mesoderm induction efficiency at day 3 of differentiation and cardiac troponin T (cTnT) was used to assess cardiomyocyte purity at day 18.

**Figure 2.**
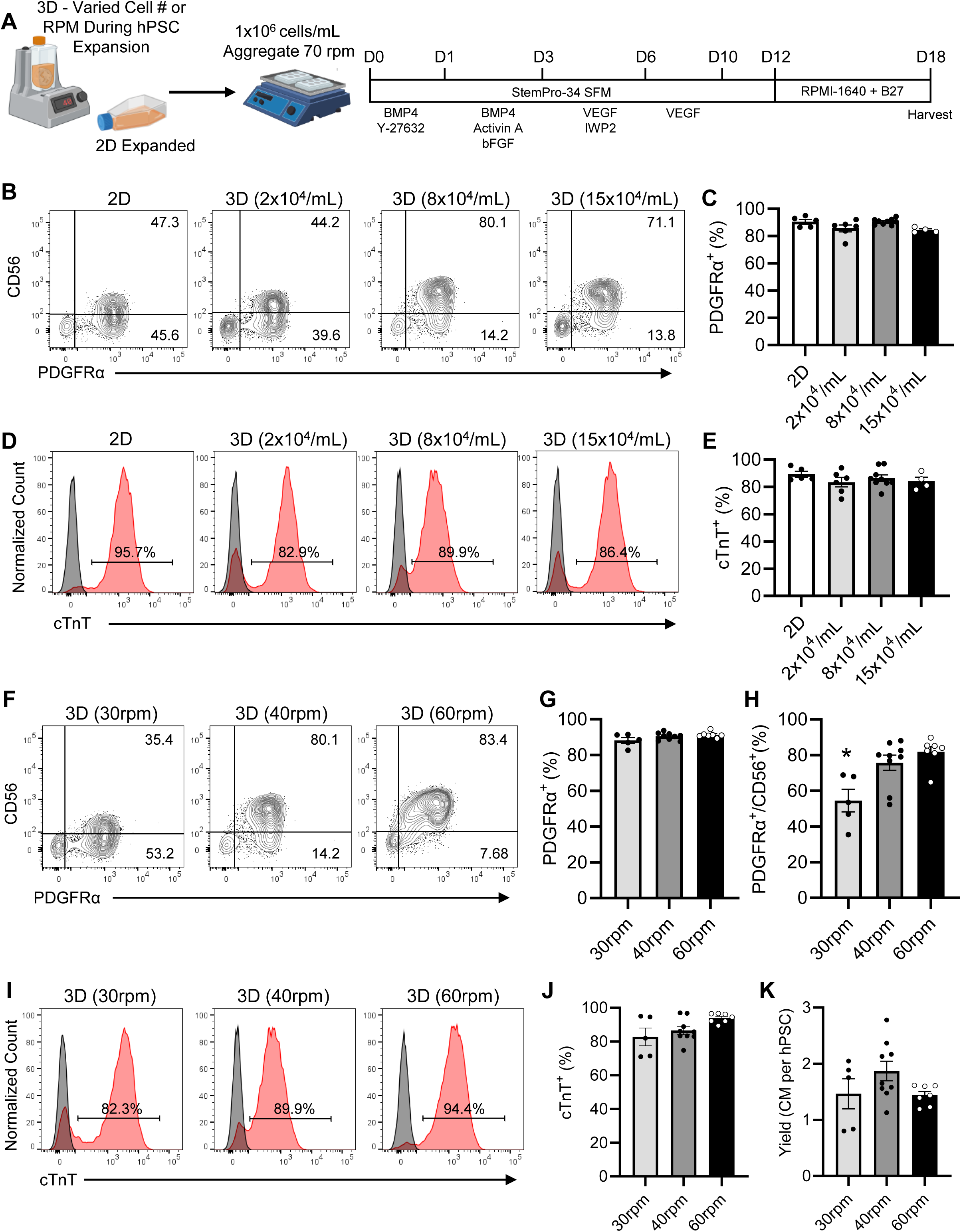
hPSCs expanded in the VWBR are compatible with a growth factor-based hPSC-CM differentiation protocol. A) Growth factor-based cardiac differentiation protocol applied to undifferentiated hPSCs previously expanded in conventional 2D culture or in VWBRs (3D). Cells were harvested on days 16-18 of differentiation for purity analysis. B) Representative flow cytometry plots for mesodermal markers (CD56 and PDGFRα) on day 3 of differentiation of hPSCs previously expanded in 2D or in VWBRs with different seeding densities; C) quantification of % PDGFRα^+^ cells by condition. D) Representative flow cytometry plots for the cardiomyocyte marker cardiac troponin T (cTnT). E) quantification of % cTnT^+^ cells by condition. F-H) Representative flow cytometry and quantification of mesodermal markers on day 3 of differentiation of cultures derived from hPSCs expanded in the VWBR at different agitation rates. I-K) Representative flow cytometry for % cTnT^+^ and yield of cTnT^+^ cells in day 16-18 cultures. *p<0.05 30 rpm vs. all other groups by one-way ANOVA followed by a Tukey post hoc. N=5-9/group

We began by testing the cardiac potential of undifferentiated hPSCs after 3D expansion in VWBRs from various initial seeding densities. Both 2D and 3D-expanded hPSCs showed comparably robust mesoderm induction and gave rise to a similar fraction of PDGFRα^+^ cells on day 3 (**Figure 2B-C**). However, the fraction of double-positive PDGFRα^+^/CD56^+^ cells was somewhat higher for hPSCs that had been expanded in VWBRs from an initial seeding density of 8x10^4^ cells/mL (**Supplementary Figure 5A**). All conditions gave rise to populations of similarly high cardiomyocyte purity (cTnT^+^) and yield when harvested on day 18 (**Figure 2D-E & Supplementary Figure 5B**). Next, we compared the cardiac potential of undifferentiated hPSCs from VWBRs operated at different agitation rates (30, 40 and 60 rpm). Here again, all conditions resulted in a similar fraction of PDGFRα^+^ cells on day 3 (**Figure 2F-G**), although the fraction of PDGFRα^+^/CD56^+^ cells was higher for hPSCs that had been exposed to higher agitation rates (**Figure 2H**). All conditions differentiated into hPSC-CMs of similar purity and yield **(Figure 2I-K)**, but there was a trend toward higher cardiomyocyte purity but lower yield for hPSCs from VWBRs operated at 60 rpm vs. 40 rpm. Finally, we compared the cardiac differentiation potential of undifferentiated hPSCs expanded in VWBRs using either StemScale or mTeSR1 medium and the hPSC expansion protocol listed in **Supplementary Figure 4A**. Surprisingly, hPSCs expanded in StemScale exhibited lower mesoderm commitment on day 3 (**Supplementary Figure 5C-D**). This did not correlate to lower cardiomyocyte purity on day 18 (**Supplementary Figure 5E**), but there was a trend toward reduced cardiomyocyte yield for undifferentiated hPSCs expanded in StemScale versus mTeSR1 (**Supplementary Figure 5F**). Thus, while hPSCs cultured in StemScale showed robust expansion and were capable of cardiac differentiation, we focused on mTeSR1 medium for subsequent experiments, given its more consistent cardiomyocyte yields.

### Determining the effect of differentiation medium composition on hPSC-CM yield

Before advancing to establish cardiac differentiation in the larger-scale VWBR system, we first used our usual 6-well plate format to test two different medium formulations during the later stages of our guided differentiation protocol. Our standard cardiac differentiation protocol involves a switch from StemPro to RPMI-B27 medium on day 12, but here we compared cardiac purity and yield obtained by this approach to a modified protocol in which the differentiating cultures were instead transitioned to diluted StemPro medium (25:75 v/v mixture of StemPro and IMDM medium, hereafter ‘SP/IMDM’) on day 6. For this, we generated undifferentiated hPSCs in VWBRs (expanded from an initial seeding density of 8x10^4^ cells/mL and operated at 40 rpm), which were then dispersed to single cells and differentiated as depicted in **Supplementary Figure 6A**. Interestingly, while hPSC-CM purity and yield were comparable between the two protocols (**Supplementary Figure 6B-D**), the fraction of hPSC-CMs expressing the committed ventricular marker MLC2v was significantly higher in cultures generated using SP/IMDM medium (**Supplementary Figure 6E-F**). Moreover, a comparison of cardiomyocyte gene expression at day 20 showed cardiac markers MYL2, MYL7, and SERCA2a were upregulated in SP/IMDM versus RPMI-B27 cultures, while HCN4 and MYH7 expression were lower (**Supplementary Figure 6G**). Given these results, differentiating cultures were transferred into SP/IMDM medium on day 6 during all subsequent experiments.

### Adapting the cardiac differentiation protocol to the 100 mL VWBR format

Having demonstrated the cardiac potential of undifferentiated hPSCs expanded in VWBRs, we next worked to test the feasibility of cardiac differentiation in the VWBR system. We began with cardiac differentiations in 100 mL VWBRs with the goal of eventual scale-up to 500 mL vessels. For this, undifferentiated hPSCs were either expanded in static 2D culture or in 3D using the VWBR, again with an initial seeding density of 8x10^4^ cells/mL. At the end of the expansion protocol, the resultant hPSCs were dispersed into single cells, placed in suspension culture in either VWBRs or 6-well plates at 1x10^6^ cells/mL, and subjected to the cardiomyocyte differentiation depicted in **Figure 3A**. We began by determining the optimal agitation rate during cell re-aggregation on day 0 of the protocol, as aggregate size is a critical factor for achieving efficient cardiogenesis (target diameter: 70-100 μm). Interestingly, while hPSCs that had been expanded in static 2D culture formed appropriately sized aggregates in VWBRs at higher speeds (40rpm), we had to slow VWBRs to 30 rpm to generate appropriately sized aggregates with hPSCs previously expanded in 3D (**Figure 3B-C**). For the subsequent differentiation runs, all 3D expanded hPSCs were aggregated at 30 rpm on day 0 of the differentiation protocol in the VWBR, and all 2D expanded cells were aggregated at 40 rpm in the VWBR.

**Figure 3.**
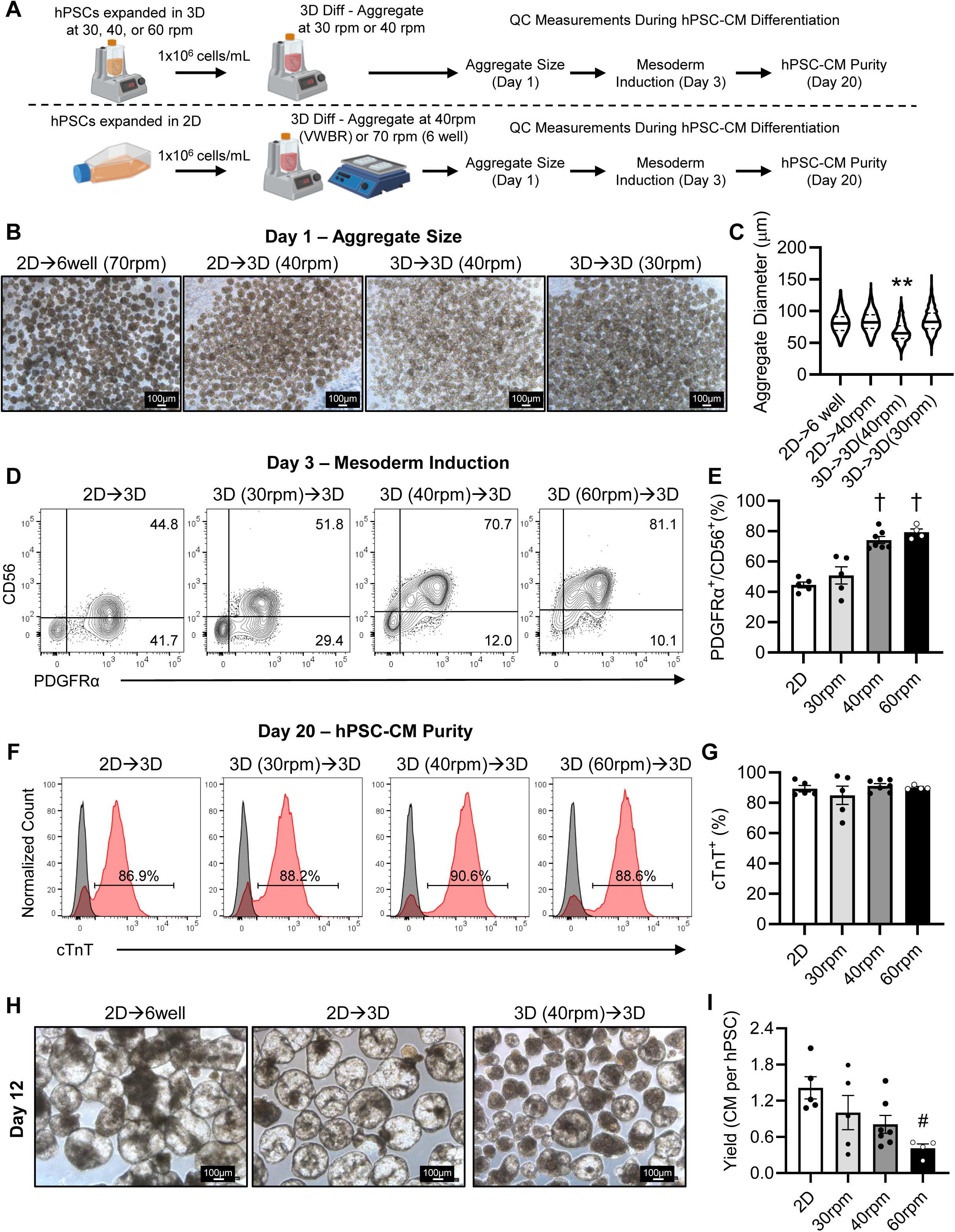
hPSC-CM differentiation in the 100 mL VWBR. A) Schematic of hPSC differentiation in 100 mL VWBRs. hPSCs were expanded in flasks (2D) or in VWBRs at different agitation rates (30, 40, or 60 rpm). To identify the optimal aggregation speed for our cardiac differentiation protocol in the VWBR, cells were aggregated at different agitation rates on D0 after inoculating the reactor at 1x10^6^ cells/mL. Cells previously expanded in 2D and aggregated in a 6-well plate at 70 rpm were used to benchmark aggregate size for differentiation. B) Representative images of aggregates on day 1 and C) distribution of aggregate sizes formed under different agitation rates. 2D→6 well(70 rpm) = hPSCs expanded in 2D and aggregated on an orbital shaker at 70 rpm, 2D→3D(40 rpm) = hPSCs expanded in 2D and aggregated in the VWBR at 40 rpm, 3D→3D(40 rpm) = hPSCs expanded in 3D and aggregated in the VWBR at 40 rpm, and 3D→3D(30 rpm) = hPSCs expanded in 3D and aggregated in the VWBR at 30 rpm. D-I) For all subsequent experiments, the agitation rate at which hPSCs were expanded in the VWBR prior to differentiation is indicated. 3D (30 rpm)→3D = hPSCs expanded in the VWBR at 30 rpm and differentiated in the VWBR, 3D (40 rpm)→3D = hPSCs expanded in the VWBR at 40 rpm and differentiated in the VWBR, 3D (60 rpm)→3D = hPSCs expanded in the VWBR at 60 rpm and differentiated in the VWBR. D) Representative plots of mesoderm commitment on day 3 of differentiation and E) quantification of PDGFRα^+^/CD56^+^ cells. F) Representative plots of hPSC-CM purity on days 18-20 and G) quantification. H) Representative images of hPSC-CM aggregate morphology on day 12. I) hPSC-CM yield on days 16-18 per starting undifferentiated hPSC. **p<0.01 3D→3D(40 rpm) vs. all other groups, ^†^p<0.01 vs. 2D and 30 rpm, ^#^p<0.05 60 rpm vs. 2D by one-way ANOVA followed by a Tukey post hoc. N=4-7/group.

We next examined how the agitation rate (30, 40, or 60 rpm) applied during undifferentiated hPSC expansion in VWBRs affected subsequent cardiac differentiation in the VWBR system. Consistent with our previous findings in 6-well plates (**Figure 2H**), hPSCs previously expanded in VWBRs at 40 or 60 rpm had a significantly higher fraction of double-positive PDGFRα^+^/CD56^+^ cells on day 3 than did hPSCs from either static 2D cultures or VWBRs operated at 30 rpm (**Figure 3D-E**), while total PDGFRα levels were similar (**Supplementary Figure 5G**). As before, hPSCs from 2D cultures and all VWBR conditions differentiated into populations of comparably pure cTnT^+^ cardiomyocytes (**Figure 3F-G**). However, while cardiomyocyte purity was similar across conditions, VWBR-expanded hPSCs tended to form smaller hPSC-CM aggregates (**Figure 3H**) and have lower cardiomyocyte yields (number of cTnT^+^ cells per starting hPSC) (**Figure 3I**) compared to hPSCs from static 2D cultures.

To overcome the lower cardiomyocyte yields of VWBR-expanded hPSCs differentiated in the VWBR system, we analyzed the early differentiation kinetics of these cells in more detail by monitoring the fraction of PDGFRα^+^ and PDGFRα^+^/CD56^+^ progenitors arising on days 2, 2.5, and 3 of differentiation (**Figure 4A**). Interestingly, when hPSCs from static 2D cultures were differentiated in the VWBR (**Figure 4A-C**), they had a time-course of mesodermal commitment similar to differentiations in 6-well plates (**Supplementary Figure 7A-C**), with a low frequency of PDGFRα^+^ and PDGFRα^+^/CD56^+^ cells on day 2 and maximal commitment by day 3. By contrast, VWBR-expanded hPSCs exhibited more rapid differentiation kinetics, reaching near-maximal levels of PDGFRα^+^ and PDGFRα^+^/CD56^+^ progenitors by day 2.5. Of note, this phenomenon was not unique to differentiations performed in the VWBR, as we found hPSCs previously expanded in the VWBR, at either 40 rpm or 60 rpm, had similarly accelerated mesodermal commitment when differentiated in 6-well plates (**Supplementary Figure 7A-C**). Given these findings of accelerated differentiation kinetics and prior reports suggesting that an earlier timing of Wnt inhibition may help maximize cardiogenesis [8], we tried varying the timing of Wnt inhibition during the differentiation protocol. Unfortunately, earlier Wnt inhibition did not enhance cardiogenesis in VWBRs; instead, we found our usual timing of IWP2 treatment on day 3 resulted in the highest fraction of cTnT^+^ cells on day 18 (**Figure 4D**).

**Figure 4.**
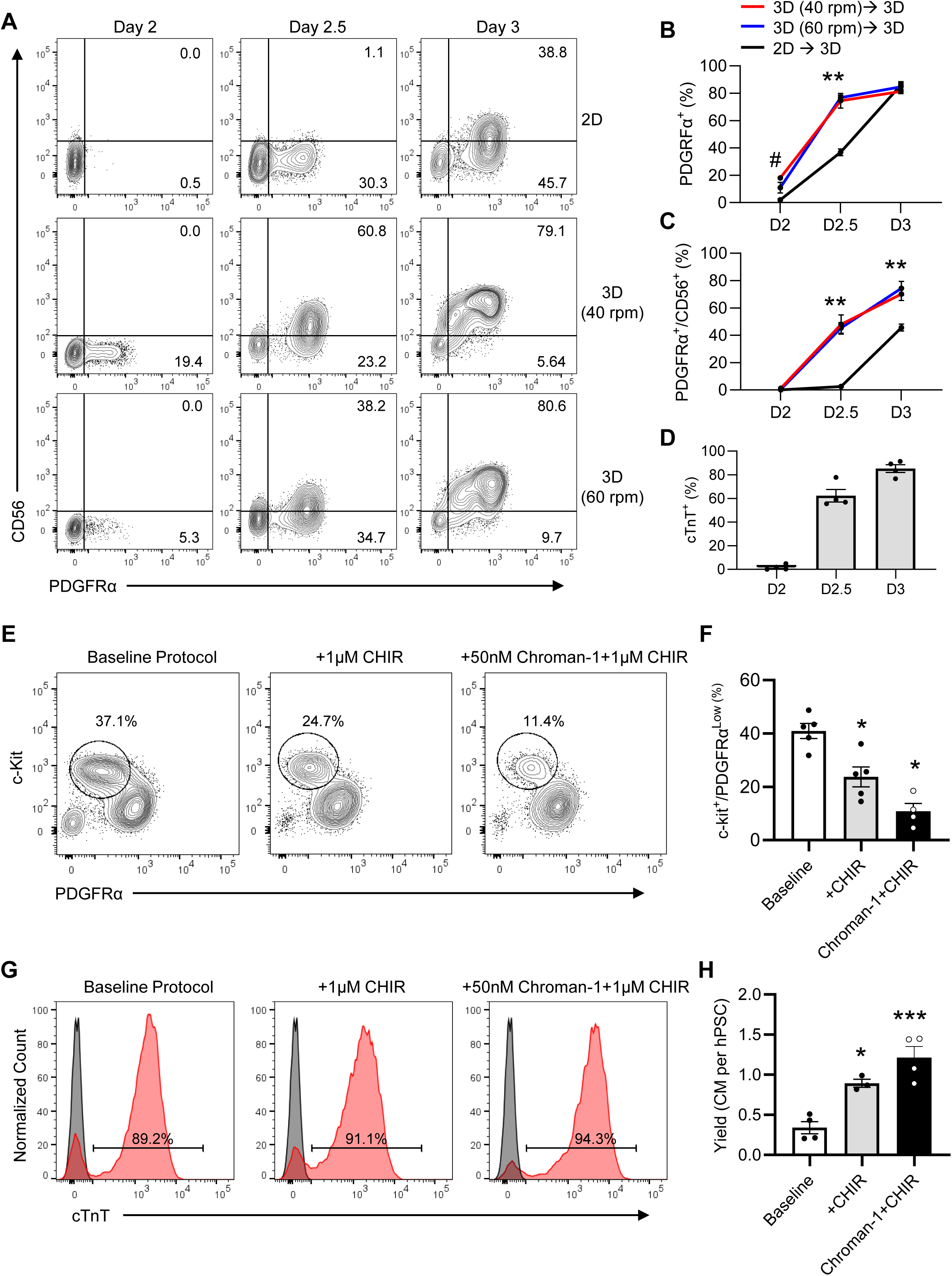
Optimization of hPSC-CM yield during differentiation in the VWBR. To assess the mesoderm commitment kinetics of hPSCs expanded in the VWBR, samples were taken from VWBRs on days 2, 2.5, and 3 of differentiation and mesodermal markers were assessed by flow cytometry. Cells cultured in conventional 2D culture and differentiated in the VWBR were used as a control. A) Representative plots assessing mesoderm commitment on days 2, 2.5 and 3 of differentiation. B) Quantification of PDGFRα^+^ and C) PDGFRα^+^/CD56^+^ cells over the same period. N=4-5/group. D) To determine whether the timing of Wnt inhibition impacts hPSC-CM purity, cells were removed from the VWBR on days 2, 2.5 and 3 and treated with the Wnt inhibitor IWP2. Cells were maintained until day 14, and hPSC-CM purity was assessed by flow cytometry. N=4/group. To assess our cell population for off-target lineage emergence, cells were assessed by flow cytometry for the presence of c-Kit+ cells on day 3. Modification of the mesodermal induction (day 1-3) stage using 1 µM CHIR and the aggregation stage (day 0-1) using 50 nM chroman-1 was also tested to assess the ability of these modifications to reduce the number of c-Kit^+^ cells on day 3. E-F) Representative plots and quantification of c-kit/PDGFRα^low^ cells on day 3 of the differentiation protocol. G) hPSC-CM purity and H) yield at day 14 of the same groups. ^#^p<0.05 40 rpm vs. 2D, **p<0.01 2D vs. all other groups by two-way ANOVA followed by Tukey post hoc. *p<0.05 vs. Baseline, and ***p<0.01 Chroman-1+ CHIR vs. Baseline by one-way ANOVA followed by a Tukey post hoc.

Next, we analyzed our VWBR differentiations for lineage contaminants, as altered cell signalling in VWBRs may promote off-target lineage formation and limit the number of cardiomyocyte progenitors. hPSCs expanded at 60 rpm in the VWBR with a seeding density of 8x10^4^ cells/mL were used for these experiments since this condition had the poorest yield (**Figure 3I**). We found that differentiation of VWBR-expanded hPSCs included a significant fraction (∼40%) of c-Kit^+^/PDGFRα^low^ cells as early as day 3 (**Figure 4E-F**). This contaminating cell population was c-Kit^+^/CXCR4^+^ and PDGFRα^-^ on day 4, suggesting off-target endodermal differentiation may be occurring (**Supplementary Figure 7D**). Interestingly, our culture conditions did not support the maintenance of these cells, as the final cell population on day 18 was comprised of >85% cTnT^+^ cardiomyocytes **(Figure 4G & Supplementary Figure 7E**). The addition of 1 µM CHIR during mesoderm induction at day 1 significantly reduced this early off-target lineage contamination and greatly enhanced hPSC-CM yield, bringing us closer to our target yield of 1 hPSC-CM per 1 input hPSC (**Figure 4E-H**).

With the goal of further improving hPSC-CM yield, we tried modifying the cell re-aggregation step (day 0) of our protocol by replacing the ROCK inhibitor Y-27632 with Chroman-1. Chroman-1 is a more potent and selective ROCK inhibitor than Y-27632, and has been reported to facilitate single-cell passaging of hPSCs and hPSC derivatives at higher viability than Y-27632 [26]. Consistent with this, we found that replacing 10 µM Y-27632 with 50nM Chroman-1 on day 0 further reduced off-target lineage commitment in VWBR cultures and significantly increased hPSC-CM yield beyond that achieved by CHIR addition alone (**Figure 4E-H & Supplementary Figure 7D-E**). Importantly, hPSC-CM purity was unaffected by these modifications (**Figure 4G and Supplementary Figure 7E**). Taken together, these refinements to our protocol (optimized VWBR settings, addition of Chroman-1 on day 0, addition of CHIR on day 1) enabled the reliable generation of >90% cTnT^+^ hPSC-CMs at final densities of >1x10^6^ cells/mL in 100 mL VWBRs.

### Generation of atrial- versus ventricular-like cardiomyocytes in VWBRs

A potential advantage of our growth factor-based differentiation protocol is that it provides controllable patterning of mesoderm subtypes during induction [15, 18, 27]. Prior work in model organisms has demonstrated that cardiomyocyte subtypes arise from distinct precursor cells associated with different waves of cardiogenesis during development, commonly referred to as the first heart field (FHF) and second heart field (SHF) [28]. By modulating the BMP4 and Activin A concentrations during the early steps of hPSC differentiation, one can generate distinct mesodermal populations that resemble FHF and SHF cells, thereby enabling the generation of different cardiomyocyte subtypes [18, 27, 29]. To determine whether undifferentiated hPSCs expanded in VWBRs retain a similar capacity to respond to varied cytokine levels and form these distinct populations, we differentiated VWBR-expanded hPSCs in 6-well plates using a range of different BMP4 and Activin A concentrations, then compared their relative flow cytometry profiles on days 3 and 4 using previously reported markers of cardiac mesoderm and SHF progenitors [18, 27]. In brief, VWBR-expanded hPSCs responded appropriately to low concentrations of BMP4 and Activin A, differentiating into cells with a SHF-like profile on day 4, including positive aldehyde dehydrogenase (ALDH) activity and low CD235a expression (**Supplementary Figure 8A-D**).

Next, we assessed whether this patterning could be reproduced during differentiation in the VWBR system and whether the resultant SHF mesoderm population could be used to generate atrial-like cardiomyocytes. For this, undifferentiated hPSCs were expanded in VWBRs and subjected to the cardiac differentiation protocol depicted in **Figure 5A**, followed by analysis of mesoderm and SHF markers on days 3 and 4. We tested induction with two different BMP4 and Activin A concentration ratios: our “standard” ventricular induction with 10 ng/mL BMP4 and 6 ng/mL Activin A (abbreviated 10B:6A), and an induction protocol expected to favor SHF specification with 4 ng/mL BMP4 and 2 ng/mL Activin A (abbreviated 4B:2A). While both the 10B:6A and 4B:2A conditions resulted in similar proportions of PDGFRα^+^ and CD56^+^ progenitors at day 3, they showed very distinct patterns of CD235a expression and ALDH activity on day 4, implying successful induction of FHF and SHF mesodermal populations, respectively (**Figure 5B-C**). Having confirmed the efficient induction of SHF mesoderm in the VWBR system, we next tested a full atrial differentiation protocol, modified from [18] that involved mesoderm induction with 4B:2A and treatment with 0.5 µM retinoic acid (RA) on day 3. Cultures resulting from this atrial differentiation protocol had a very different morphology relative to those from our usual ventricular differentiation protocol (10B:6A and no RA). Cell aggregates from the atrial protocol appeared smaller and denser, while aggregates from the ventricular protocol were larger with a more cystic structure (**Figure 5D**). Cardiac marker expression on day 20 was consistent with the generation of different cardiac subtypes, with the atrial protocol generating primarily MLC2a^+^ cells and the ventricular protocol generating both MLC2v^+^ and MLC2a^+^ cells, as expected for immature ventricular cardiomyocytes [30–32] (**Figure 5E-H**). Comparison of gene expression signatures further supports successful production of distinct cardiomyocyte subtypes [18], as cells from the atrial protocol strongly expressed TBX5, NR2F2, and KCNJ3, while cells from the ventricular protocol strongly expressed MYL2 and IRX4 (**Figure 5I**).

**Figure 5.**
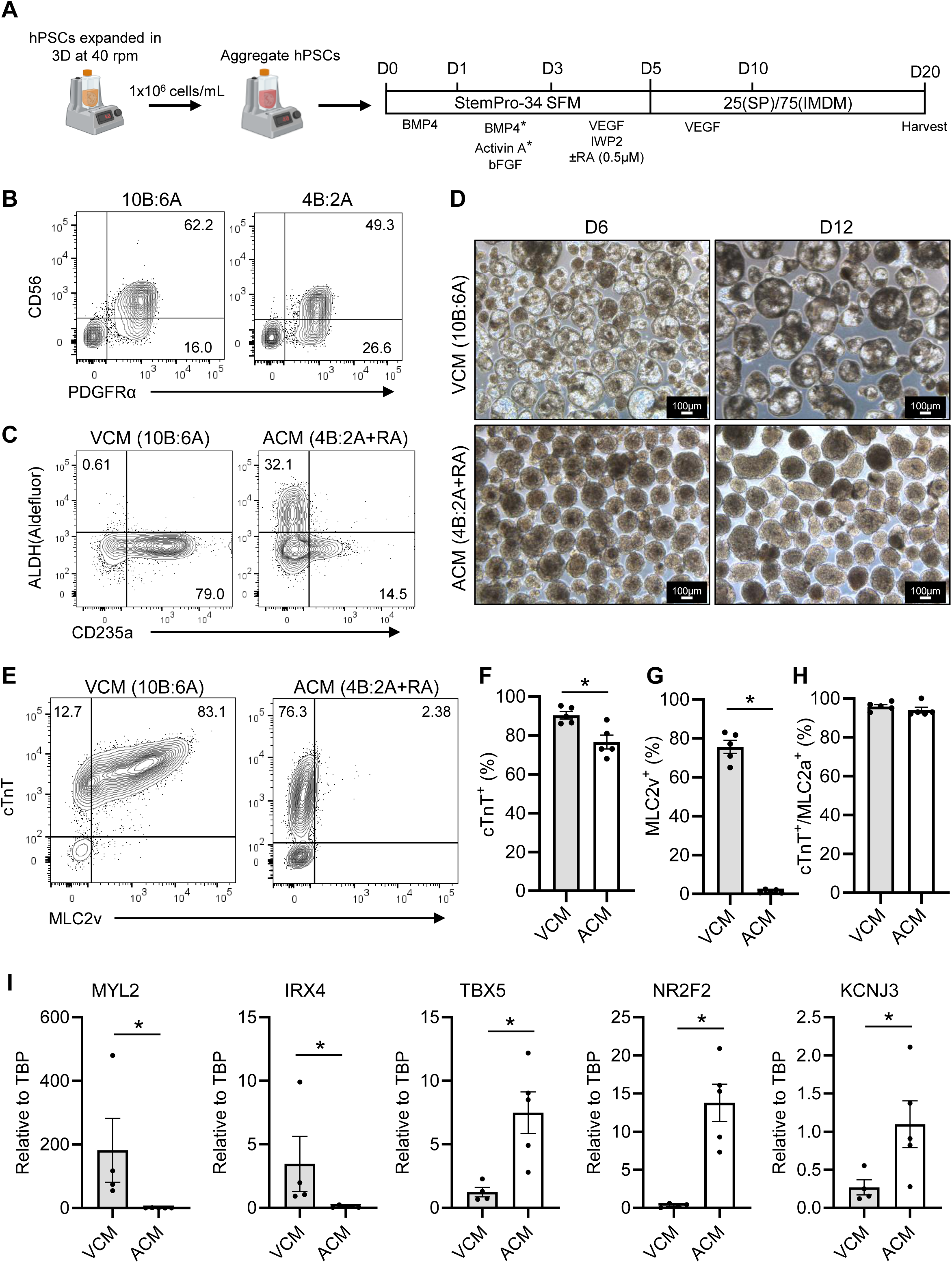
Differentiation of ventricular and atrial CMs in 100 mL VWBRs. A) Schematic of the protocols used to produce either ventricular CMs or atrial CMs. The ventricular differentiation protocol (referred to as 10B:6A) involves mesoderm induction with 10 ng/mL BMP4 and 6 ng/mL activin A from days 1-3, while the atrial protocol (referred to as 4B:2A) involves mesoderm induction using 4 ng/mL BMP4 and 2ng/mL activin A as well as addition of 0.5 µM retinoic acid (RA) from day 3-5. Cells were assessed for mesoderm induction efficiency using PDGFRα and CD56 on day 3. On day 4, ALDH activity and CD235a expression were used to identify second heart field (SHF) and first heart field (FHF) populations. B) Representative mesoderm profile at day 3 and C) day 4 of the differentiation protocol. D) Representative images of aggregate morphology on days 6 and 12 following differentiation with either the ventricular or atrial protocol. E) Representative flow cytometry plots for pan-cardiac marker cTnT and ventricular marker MLC2v in day 20 cultures generated with each differentiation protocol. Quantification of the percentage of cells in each condition expressing F) cTnT^+^, G) MLC2v, and H) MLC2a. I) Expression of ventricular or atrial genes at day 20 of differentiation. *p<0.05 by two-tailed Student’s t-test.

### Generation of hPSC-CMs in 500 mL VWBRs

After successfully adapting our cardiac differentiation protocol to 100mL VWBRs, we next sought to scale up our protocol to 500 mL VWBRs with a target yield of >500x10^6^ hPSC-CMs per reactor vessel. Because we needed to scale up undifferentiated hPSC production similarly, we began by assessing our hPSC expansion in 500 mL VWBRs. Fortunately, our hPSC expansion protocol translated from 100 mL to 500 mL reactors, and we identified a protocol by which 500 mL VWBRs were seeded with hPSCs at an initial density of 8x10^4^/mL and operated at 30 rpm (**Supplementary Figure 9A**). hPSC expansion over 5 days using this protocol sufficed to reach our target of >1x10^6^ cells/mL at >90% viability (**Supplementary Figure 9B**). Importantly, aggregate diameter, glucose consumption, lactate production, and pluripotency marker expression were similar to those previously described for 100 mL VWBRs (**Supplementary Figure 9C-G**).

Next, we upscaled hPSC-CM differentiation to the 500 mL VWBR format using hPSCs expanded in the 500 mL VWBR and the optimized protocol shown in **Figure 6A**. While we found that cell re-aggregation on day 0 was generally more variable in 500mL VWBRs than with the 100 mL VWBR, a reasonable distribution of aggregate sizes could be achieved using an agitation rate of 20 rpm (**Figure 6B**). 500 mL VWBR expanded hPSCs exhibited robust mesoderm induction (**Figure 6C**), and high levels of CD56 expression on day 3 of differentiation (**Figure 6D**), consistent with our observations in the 100mL VWBR (**Figure 3E**). 500 mL VWBR expanded hPSC populations yielded high purity cTnT^+^ hPSC-CMs on day 18, corresponding to >1 hPSC-CM per 1 input hPSC (**Figure 6E-F**). Interestingly, across 100 mL and 500 mL VWBRs we found that MLC2v^+^ cell frequency at day 20 negatively correlated with hPSC-CM yield (**Figure 6G**), suggesting that accelerated acquisition of the ventricular specification marker may be associated with less cell proliferation. Next, we further characterized our hPSC-CM cell product using several methodologies. Immunofluorescent staining of sectioned aggregates for cardiac troponin T confirmed high-purity hPSC-CMs within the aggregates, consistent with our flow cytometry data (**Figure 6H**). Quantification of the force-generating capacity of VWBR produced hPSC-CMs using the “Biowire” engineered heart tissue preparation [22] demonstrated that hPSC-CMs generated a mean force of > 1 mN/mm^2^ (**Figure 6I**), a value comparable to that reported for hPSC-CMs from other bioreactor-based protocols [8, 33]. Electrophysiological characterization of hPSC-CM monolayer action potentials and intercellular calcium transients by optical mapping demonstrated that VWBR produced hPSC-CMs exhibited the expected ventricular action potential morphology (**Figure 6J**), with an average spontaneous beat rate of ∼34 beats per minute (**Figure 6K**). Moreover, hPSC-CM monolayers exhibited the expected action potential and calcium transient responses to a progressively increasing pacing frequency from 1 to 2 Hz, evidenced by a smaller calcium transient amplitude (**Figure 6L**), shorter action potential and calcium transient durations (**Supplementary Figure 10A-B**), and a faster calcium transient time to peak (**Supplementary Figure 10C**). Lastly, we examined MLC2v expression of VWBR produced hPSC-CMs during prolonged culture. Although hPSC-CMs generated using our 500 mL VWBR protocol had variable expression of the ventricular marker MLC2v on day 20, the percentage of MLC2v^+^ cells increased with longer culture duration, either in suspension or after re-plating onto 2D substrates (**Figure 6M-N**), as would be expected for ventricular-specified hPSC-CMs. Importantly, high cTnT^+^ purity was maintained during this extended culture period (**Supplementary Figure 10D**).

**Figure 6.**
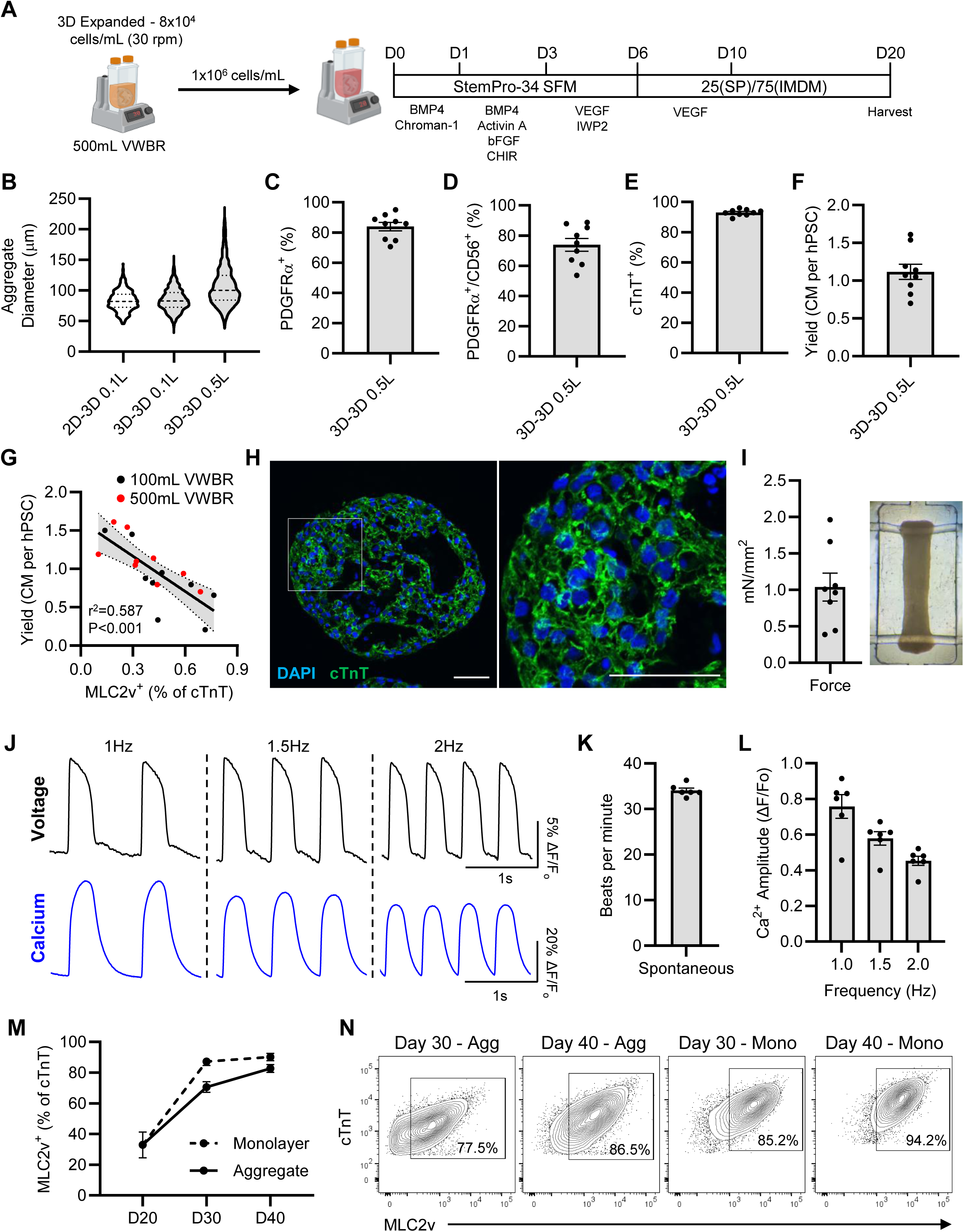
hPSC-CM production in 500 mL VWBRs. A) Protocol used to differentiate hPSCs in the 500 mL VWBR. B) hPSCs expanded in 500mL VWBRs were aggregated at 20 rpm, and the resultant aggregates were compared to our optimized conditions in the 100 mL VWBR. B) Quantification of PDGFRα^+^ and C) PDGFRα^+^/CD56^+^ cells on day 3 of differentiation in the 500 mL VWBR. Cells were maintained until day 20 and assessed for cardiomyocyte purity and yield. E) Quantification of % cTnT^+^ cells and F) cardiomyocyte yield at day 20. G) Correlation of MLC2v^+^ cell frequency and hPSC-CM yield on day 20 of differentiation, indicating a significant negative correlation between yield and MLC2v^+^ cell frequency. H) Representative cTnT staining of an hPSC-CM aggregate harvested at day 20 from a 500 mL VWBR. Scale bar, 50 µm. I) Force generation of biowires formed from different batches of day 20 hPSC-CMs produced in the 500mL VWBR. J) Representative optical action potentials and intercellular calcium transients obtained from hPSC-CM monolayers paced at 1, 1.5, and 2 Hz. K) Spontaneous beat rate of hPSC-CM monolayers formed from 500 mL VWBR produced hPSC-CMs. L) Ca^2+^ transient amplitude of hPSC-CM monolayers during electrical pacing at 1, 1.5, and 2 Hz. M) Quantification of cTnT^+^/MLC2v^+^ cell frequency at days 20, 30, and 40 of differentiation and N) Representative flow cytometry plots. Cells were maintained as aggregates or monolayers from day 20 until day 40 following harvest from the 500 mL VWBR at day 20.

Lastly, to assess the use of our VWBR produced hPSC-CMs as a cell therapy product, we transplanted 70x10^6^ hPSC-CMs into the infarcted guinea pig myocardium using our established MI model [17]. VWBR produced hPSC-CMs maintained high viability after cryo-recovery, with a mean cell viability of 78% (**Supplementary Figure 10E**). hPSC-CMs were transplanted into the infarcted myocardium 10 days after injury, and graft size was assessed 14 days after transplantation via immuno-staining of α-actinin and the human nuclear marker Ku80 (**Figure 7A**). VWBR produced hPSC-CMs exhibited robust engraftment (**Figure 7B**), with a mean graft size of 15% of the scar area (**Figure 7C**), a level of engraftment comparable to previously reported graft sizes for 2D-produced hPSC-CMs [17, 34].

**Figure 7.**
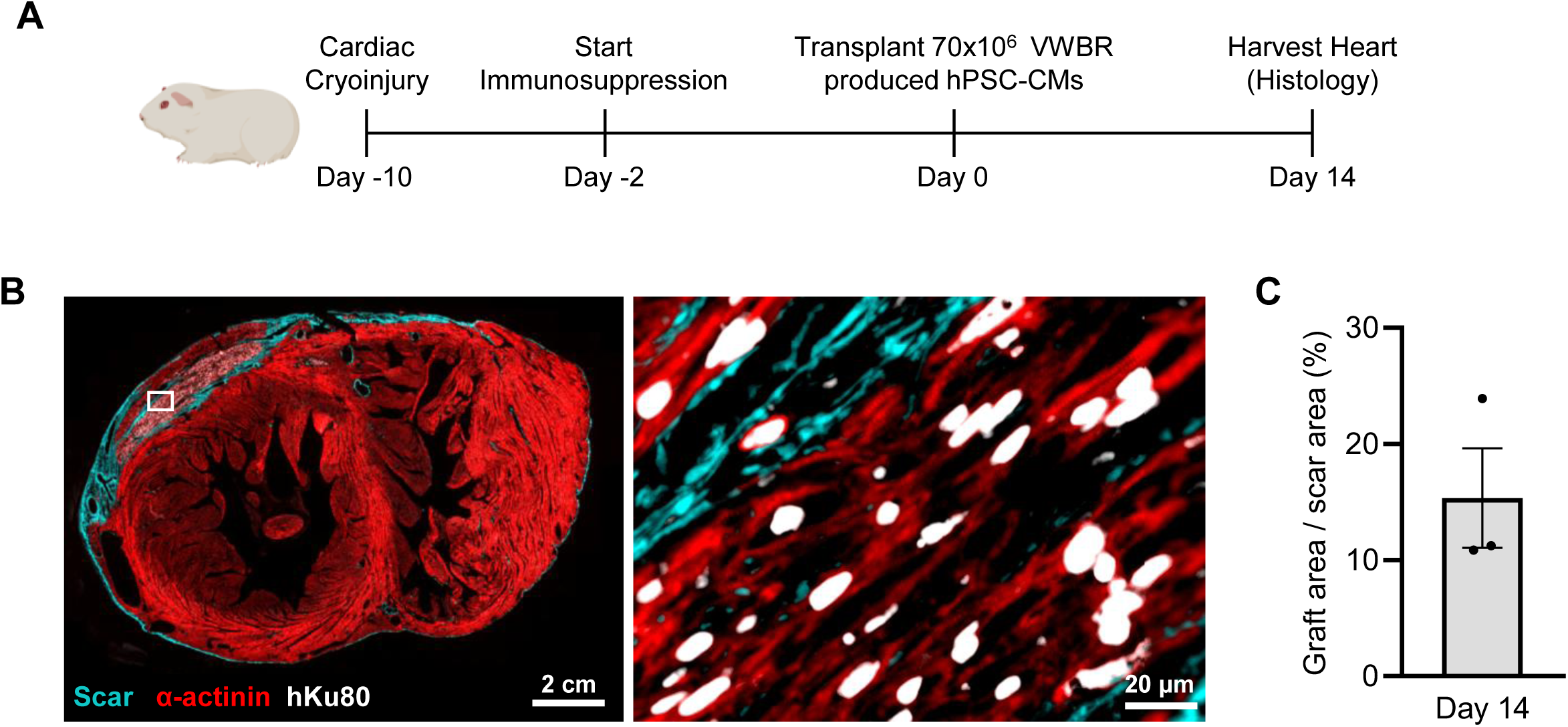
Transplantation of VWBR produced hPSC-CMs in the infarcted Guinea pig model. A) Schematic illustrating the timing of cryo-injury, immunosuppression initiation, hPSC-CM transplantation, and tissue harvest. B) Representative cross-sectional images of a Guinea pig heart 14 days following hPSC-CM transplantation (left) and inset (right) showing human cardiomyocyte graft tissue staining positive for the human nuclear marker Ku80 and the muscle marker α-actinin. C) Quantification of graft size at day 14 post-transplantation.

## Discussion

In this study, we present a scalable cardiomyocyte differentiation protocol capable of producing high purity and yields of hPSC-CMs using the VWBR system. At the 500 mL scale, this protocol yielded hPSC-CM populations with a mean cardiomyocyte purity of 93% and yield of 1.11x10^6^ hPSC-CMs/mL. To achieve this, we systematically optimized several critical parameters, including hPSC expansion, embryoid body aggregation, and mesoderm induction. We found that hPSCs expanded in the VWBR exhibited altered commitment kinetics and greater formation of cKit^+^/CXCR4^+^/PDGFRα^-^ cells during differentiation, which correlated with lower yields compared to hPSCs expanded in conventional 2D culture. Modification of the aggregation and mesoderm induction steps reduced the presence of this cell population and improved hPSC-CM yields, resulting in final yields of >1x10^6^ hPSC-CMs/mL in 100 mL and 500 mL VWBRs. Moreover, we demonstrate that our bioreactor cardiomyocyte differentiation protocols can be easily adapted to generate either atrial or ventricular cardiomyocytes. Lastly, we show that ventricular hPSC-CMs generated using our optimized protocols in the 500 mL VWBR robustly engraft following transplantation into the infarcted guinea pig myocardium, supporting their use in cell therapy applications.

The protocol described here differs from other previously reported bioreactor-based cardiac differentiation protocols [10, 12, 35, 36] in two important ways: the use of the VWBR system as an all-in-one hPSC expansion and differentiation platform, and the use of a growth factor-based hPSC-CM differentiation protocol. To our knowledge, this is the first protocol to employ the VWBR platform for both hPSC expansion and subsequent hPSC-CM differentiation. VWBRs are commercially available, single-use bioreactors that have a number of advantages relative to spinner flasks and stirred-tank reactors, including more homogenous mixing over a lower shear stress range, making VWBRs particularly well-suited for hPSC expansion [5]. For example, Kallos and colleagues demonstrated that hPSC aggregates in VWBRs had a more consistent size distribution than aggregates cultured in spinner flasks [5]. This is important because increasing aggregate size can limit the diffusion of nutrients and removal of waste, thereby creating heterogeneity in the resulting cell population [37]. hPSCs have been successfully expanded using other bioreactor formats [13, 38], but shear stress is known to adversely affect PSC viability and endogenous signalling [39–41]. Our findings here demonstrate that hPSCs can be successfully expanded over a range of seeding densities and agitation rates, highlighting the flexibility of the VWBR system. While the goal of this study was to provide sufficient starting material for subsequent hPSC-CM differentiation and not to develop high-density hPSC culture protocols, we identified culture conditions that reliably resulted in >1x10^6^ hPSCs/mL. This is in contrast with previously reported bioreactor protocols which optimized 3D hPSC-CM differentiation using hPSCs expanded in conventional 2D culture [12, 36].

The second way in which our protocol here differs from previously reported bioreactor-based cardiac differentiation methods is in its use of growth factors (BMP4, Activin A, and bFGF) to induce mesoderm and cardiomyocyte differentiation when used in combination with a Wnt inhibitor (e.g. IWP2). By contrast, all previous reported bioreactor-based protocols are based on the GiWi protocol established by Sean Palecek’s lab [42], which exclusively relies on small-molecule activators of Wnt signalling (e.g. CHIR99021) for mesoderm induction [8, 12, 33, 35]. Such protocols rely more heavily on endogenous hPSC signalling to facilitate cardiomyocyte specification, which is ultimately influenced by individual cell line characteristics and culture conditions. While small molecules do offer advantages in terms of cost and reduced batch-to-batch variation, recent studies have suggested that growth factor-based differentiation enables finer control over cardiomyocyte differentiation [14, 15]. For example, hPSC-CM differentiation as embryoid bodies using a growth factor-based protocol produced cardiomyocytes with lower Ki67^+^ expression, improved structural morphology, improved connexin-43 localization, and greater mitochondria density compared to hPSC-CMs produced using the conventional 2D GiWi protocol [14]. Additionally, by incorporating developmentally relevant growth factors, one can generate mesoderm subtypes that resemble FHF and SHF progenitors [27]; these progenitors can be further guided into the different cardiomyocyte subtypes found throughout the heart. hPSC-CMs produced using our ventricular protocol exhibited >80% MLC2v expression by day 40 of differentiation, supporting successful ventricular cardiomyocyte differentiation using our protocol presented here. This contrasts with small molecule-based monolayer protocols, which reported lower MLC2v levels at comparable time points [32]. Furthermore, our yield and cardiomyocyte purity achieved here are similar to previously published bioreactor protocols, which reported yields from 0.4x10^6^ -1.5x10^6^ hPSC-CMs/mL with a mean purity of >85% cTnT^+^, but more often >90% [8, 9, 12, 13, 33, 35, 43]. The range in yield is a function of several factors, some of which include the differentiation protocol, cell line, and culture vessel format.

One interesting observation from this study was that undifferentiated hPSCs from 2D cultures initially differentiated into cardiomyocytes in VWBRs much more efficiently than did undifferentiated hPSCs expanded in the VWBR. Likely underlying this, we found that the latter formed a larger fraction of c-Kit^+^/CXCR4^+^/PDGFRα^-^ cells on day 4, suggestive of biased endodermal differentiation. It is well established that the culture of hPSCs in 3D as aggregates affects key cell signalling pathways relative to 2D cultured cells, including Wnt activity [44, 45], TGFβ/Nodal signalling [46], FGF2 expression [47], and AMPK signalling [13]. For example, Branco and colleagues found that hPSCs cultured as aggregates in Aggrewells had greater expression of Nodal and the Nodal-responsive genes Lefty1 and Foxa2 relative to 2D cultures, implying that 3D culture can alter this signalling axis even in the absence of shear stress or other bioreactor-related cues [46]. Additionally, shear stress within bioreactors can also impact β-catenin activity and the expression of key pluripotency regulators [48], providing another route by which 3D hPSC culture may alter hPSC differentiation. Given the impact 3D hPSC culture has on key signalling pathways involved in primitive streak induction, mesoderm formation, and cardiomyocyte differentiation, further work is needed to understand the culture conditions that impact these key signalling axes to better control cell differentiation in 3D within bioreactors.

It is important to note that further refinement of our protocol is needed to meet GMP standards for cell production. This includes working towards a fully closed system and the use of GMP-grade reagents and media. Moreover, while the 500 mL scale is likely sufficient to meet the demand of an academic lab, further upscaling of the cell manufacturing process is needed to satisfy the scales required for industry applications. While out of scope for this study, it is important to note that the VWBR system can be scaled using the already commercially available 3, 15, and 80 L systems [49]. Increased culture scale will also require the implementation of established biomanufacturing strategies to limit batch-to-batch variability and improve cell yields. This includes incorporating constant perfusion and feedback systems to control cell culture media glucose, dissolved oxygen, and pH [10, 38, 50], as well as automating media exchanges.

## Conclusion

In conclusion, this work describes a scalable protocol for producing hPSCs and hPSC-CMs using the VWBR system. Implementation of growth factor-based differentiation protocols in the VWBR is expected to enable the production of a diverse range of cell types, including all cardiomyocyte subtypes and non-myocyte populations. We predict that this novel VWBR-based protocol will greatly enable cardiac cell therapy and tissue engineering applications, as well as help provide the cell quantities needed for in vitro disease modelling and drug discovery platforms.

## Abbreviations

ACM: Atrial-Like Cardiomyocyte
ALDH: Aldehyde dehydrogenase
BMP4: Bone morphogenetic protein 4
CM: Cardiomyocyte
cTnT: Cardiac troponin T
bFGF: Basic fibroblast growth factor
FHF: First heart field
hPSC: Human pluripotent stem cell
hPSC-CM: Human pluripotent stem cell-derived cardiomyocytes
MLC2a: Myosin light chain 2a
MLC2v: Myosin light chain 2v
SHF: Second heart field
STB: Stirred Tank Bioreactor
VLCM: Ventricular-like cardiomyocyte
VEGF: Vascular endothelial growth factor
VWBR: Vertical wheel bioreactor

## Acknowledgements

The authors thank Dr. Evan Miller (University of California-Berkeley) for kindly providing the BeRST1 voltage dye. We thank Dr. Cristine Reitz for critical review of the manuscript.

## Funding

This work was supported by the Canada Research Chairs Program (CRC-2020-00245) and the Government of Canada’s New Frontiers in Research Fund (NFRFT-2022-00447) to M.A.L

## Author Contributions

FJA and MAL conceived the study. FJA and MAL designed experiments. All authors performed experiments. FJA and MAL wrote the manuscript with input from all authors. All authors approved the final version of the manuscript.

## Ethics approval and consent to participate

Research ethics board (REB) and stem cell oversight committee approval were obtained for this work involving human pluripotent stem cells.

## Consent for publication

Not applicable.

## Competing interests

MAL and GK are co-founders and paid consultants for Bluerock Therapeutics.

## Supplementary Material

**Supplementary Figure 1.**
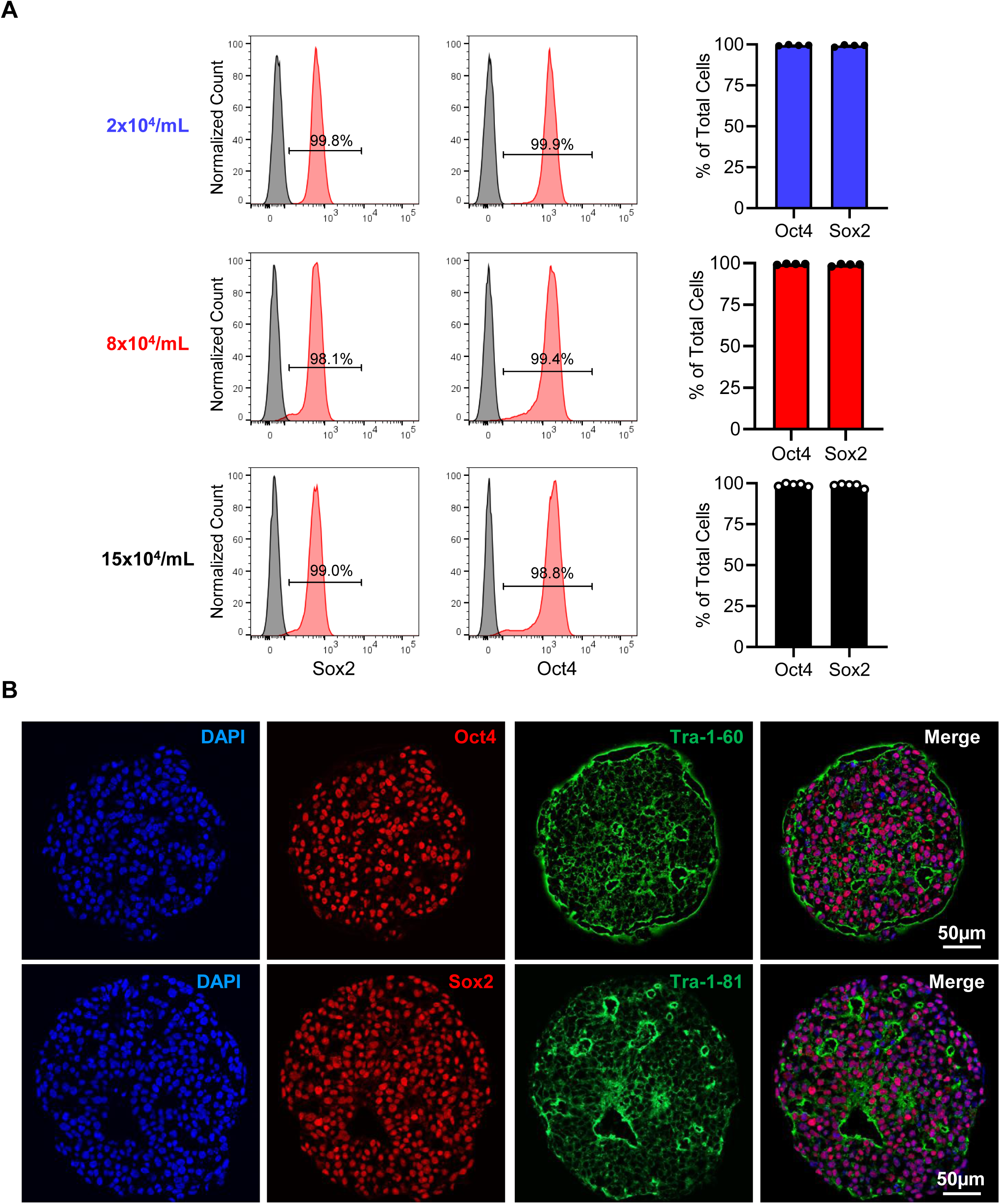
hPSC expansion in the VWBR does not impact pluripotency. A) Histogram plots and quantification of Oct4 and Sox2 expression in hPSCs expanded in the VWBR with different seeding densities. Conditions correspond to those in Figure 1. B) Representative immunofluorescent staining for Oct4 and Tra-1-60 (top) and Sox2 and Tra-1-81 (bottom) in hPSC aggregates harvested from VWBRs seeded with 8x10^4^ cells/mL and agitation rate set to 40 rpm.

**Supplementary Figure 2.**
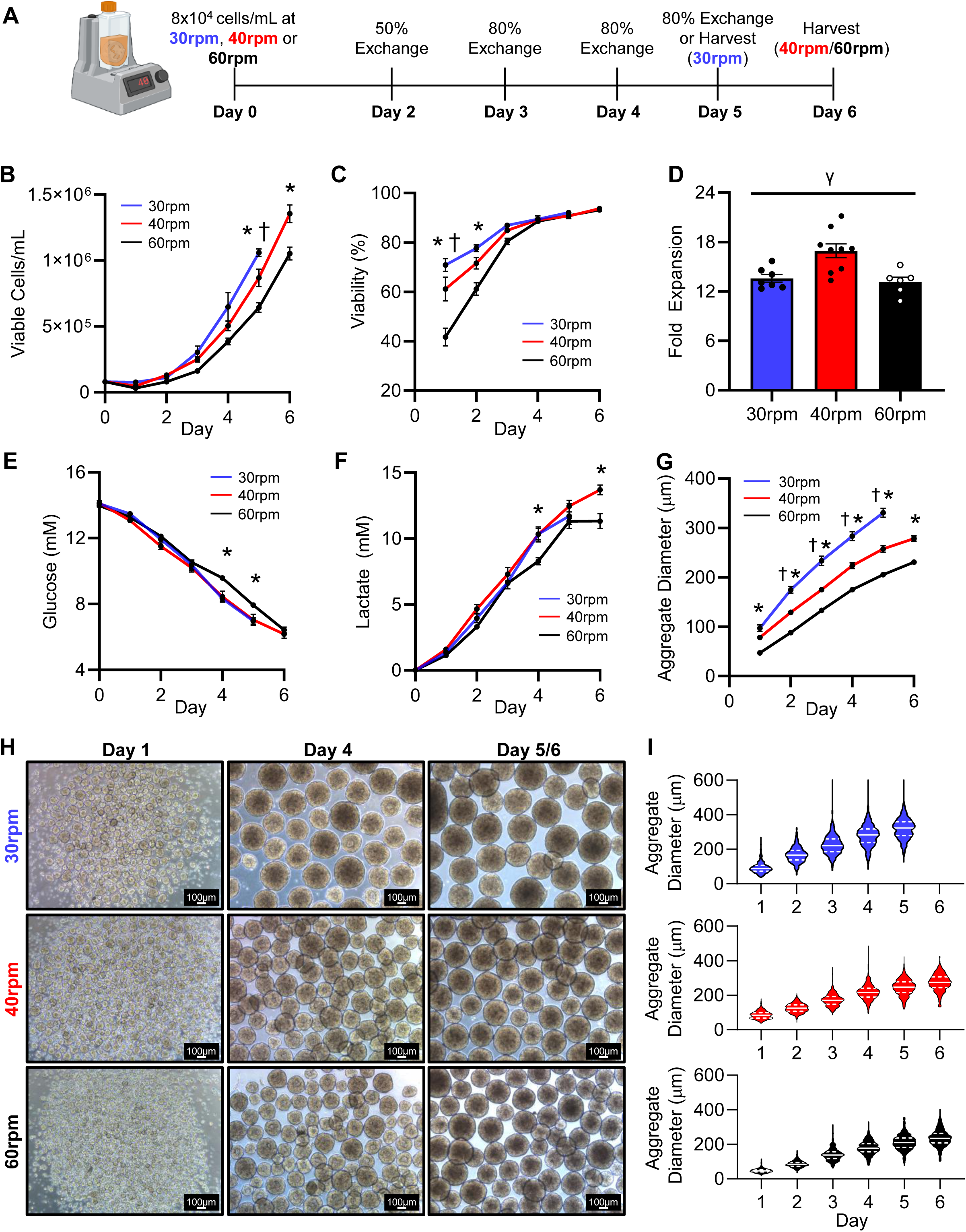
Agitation rate impacts hPSC expansion in the 100 mL VWBR. A) Schematic illustrating the strategy for media exchanges and cell harvesting for hPSC cultures maintained at different agitation rates (30, 40, or 60 rpm). Bioreactors were inoculated with 8x10^4^ cells/mL and cultured at different agitation rates for 5 or 6 days, depending on the agitation rate. B) Number of viable cells/mL and C) % viability during the expansion period. D) Fold expansion of hPSCs calculated by cell number at harvest/starting cell number on day 0. E) Media glucose and F) lactate levels, as well as G) mean aggregate diameter during hPSC expansion. H) Representative aggregate images, and I) aggregate diameter distribution during expansion in the VWBR. †p≤0.05 30 rpm vs. all other groups, *p≤0.05 60 rpm vs. all other groups by two-way ANOVA followed by a Tukey post hoc. ^γ^p≤0.05 40 rpm vs. all other groups by one-way ANOVA followed by a Tukey post hoc. N=6-9/group.

**Supplementary Figure 3.**
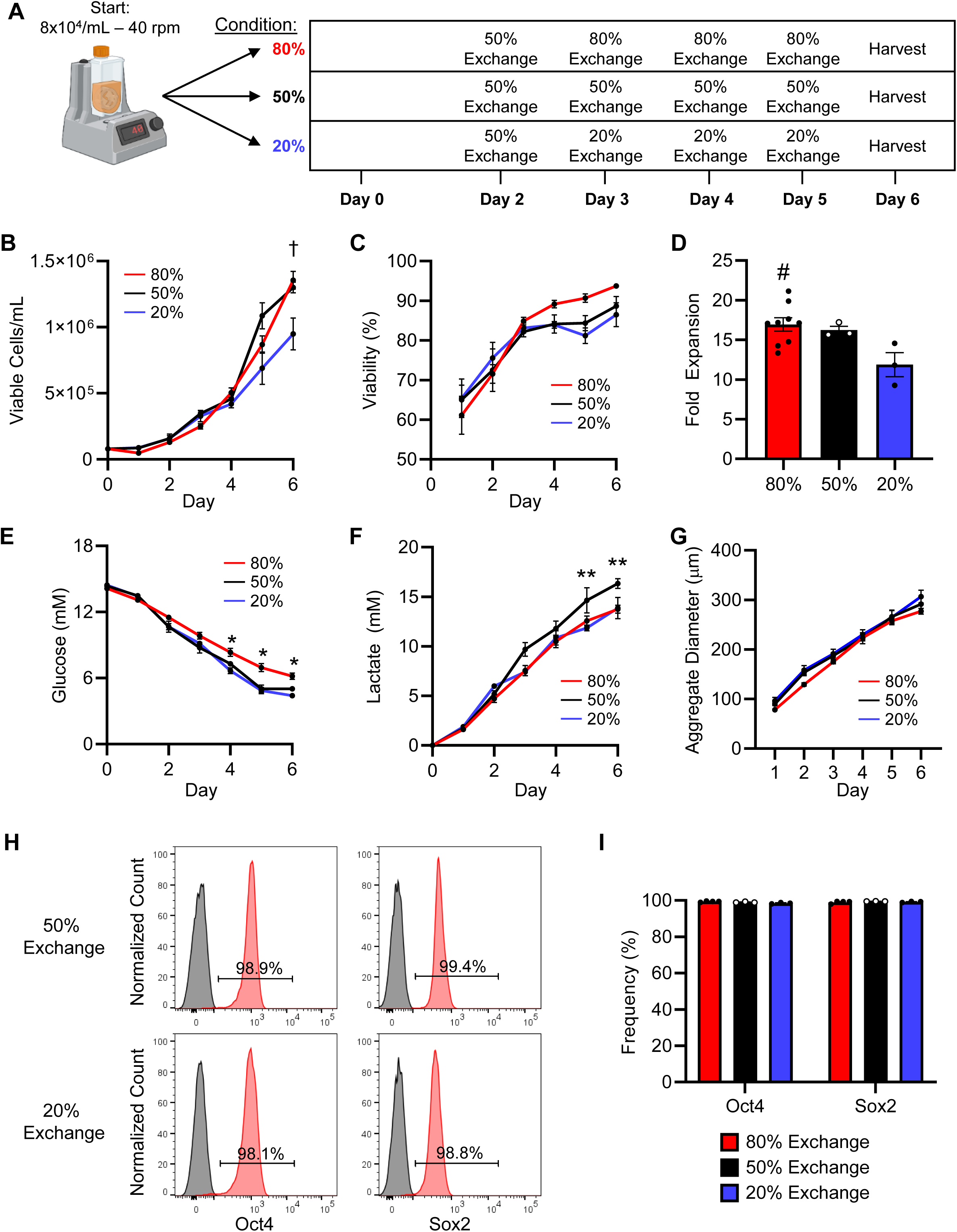
The impact of media exchange on hPSC expansion in 100mL VWBRs. A) Media exchange schedules tested for hPSC expansion in the VWBR with a seeding density of 8x10^4^ cells/mL and an agitation rate of 40 rpm. B-G: parameters for hPSCs expanded under each of the three conditions, including viable cells/mL (B), % cell viability (C), fold expansion (D), glucose (E), lactate (F), and mean aggregate diameter (G). H) Histogram plots and I) quantification of Oct4 and Sox2 expression in hPSCs following expansion in the VWBR. ^†^p<0.05 20% vs. all other groups by two-way ANOVA followed by a Tukey post hoc. #p<0.05 80% vs 20% by one-way ANOVA followed by a Tukey post hoc. *p<0.05 80% vs all other groups, and **p<0.05 50% vs. all other groups by a two-way ANOVA followed by a Tukey post hoc.

**Supplementary Figure 4.**
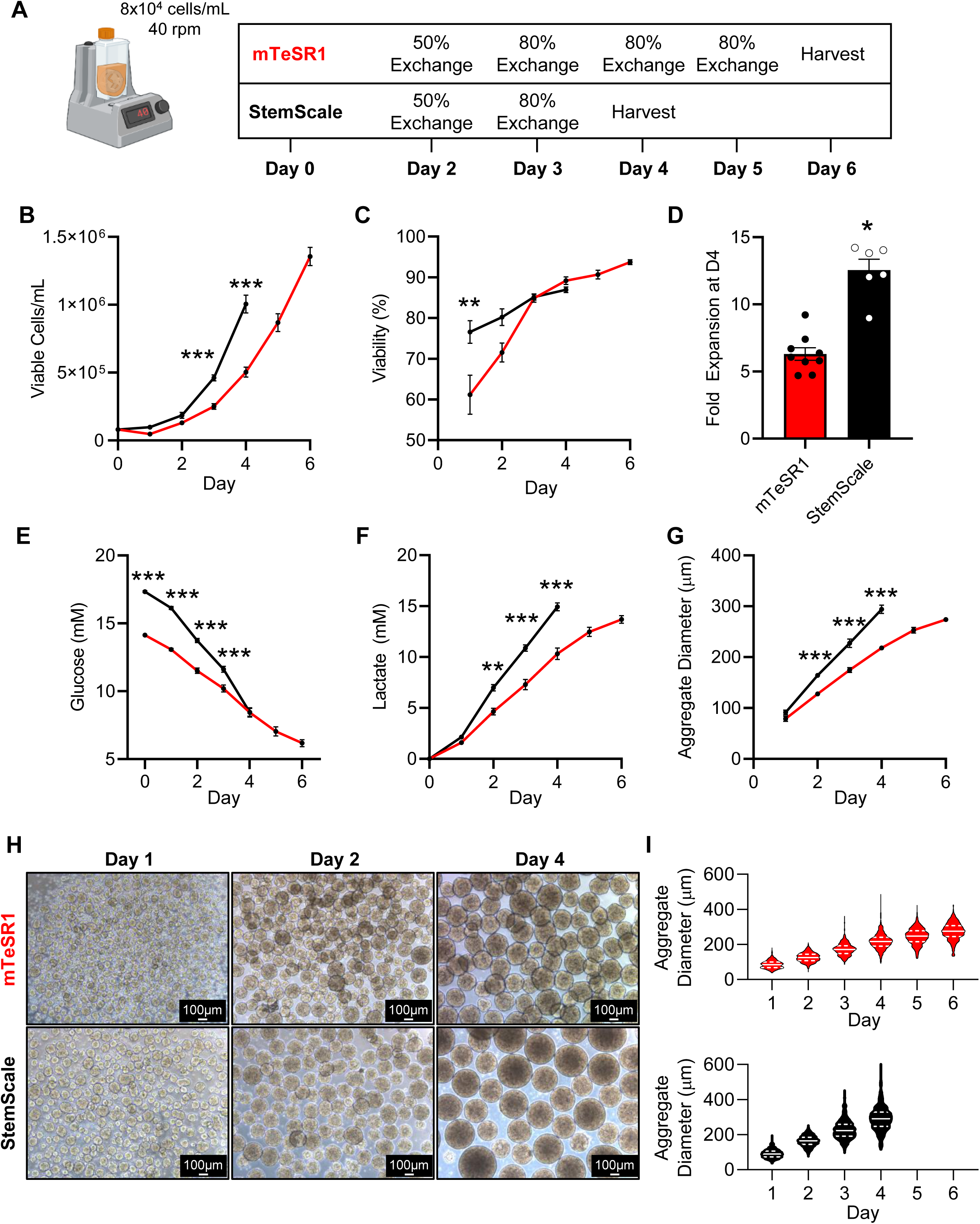
hPSC expansion in StemScale media. A) Protocol used for hPSC expansion in mTeSR1 or StemScale medium using the 100mL VWBR. B-G: parameters for hPSCs expanded in each medium, including viable cells/mL (B), % cell viability (C), fold-expansion (D), glucose (E), lactate (F), and aggregate diameter (G). H) Representative images of hPSC aggregates in each medium by day, and I) aggregate diameter distribution over the expansion period. *p<0.05, **p<0.01, ***p<0.001 StemScale vs. mTeSR1 by two-way ANOVA followed by a Tukey post hoc or Student’s t-test where applicable.

**Supplementary Figure 5.**
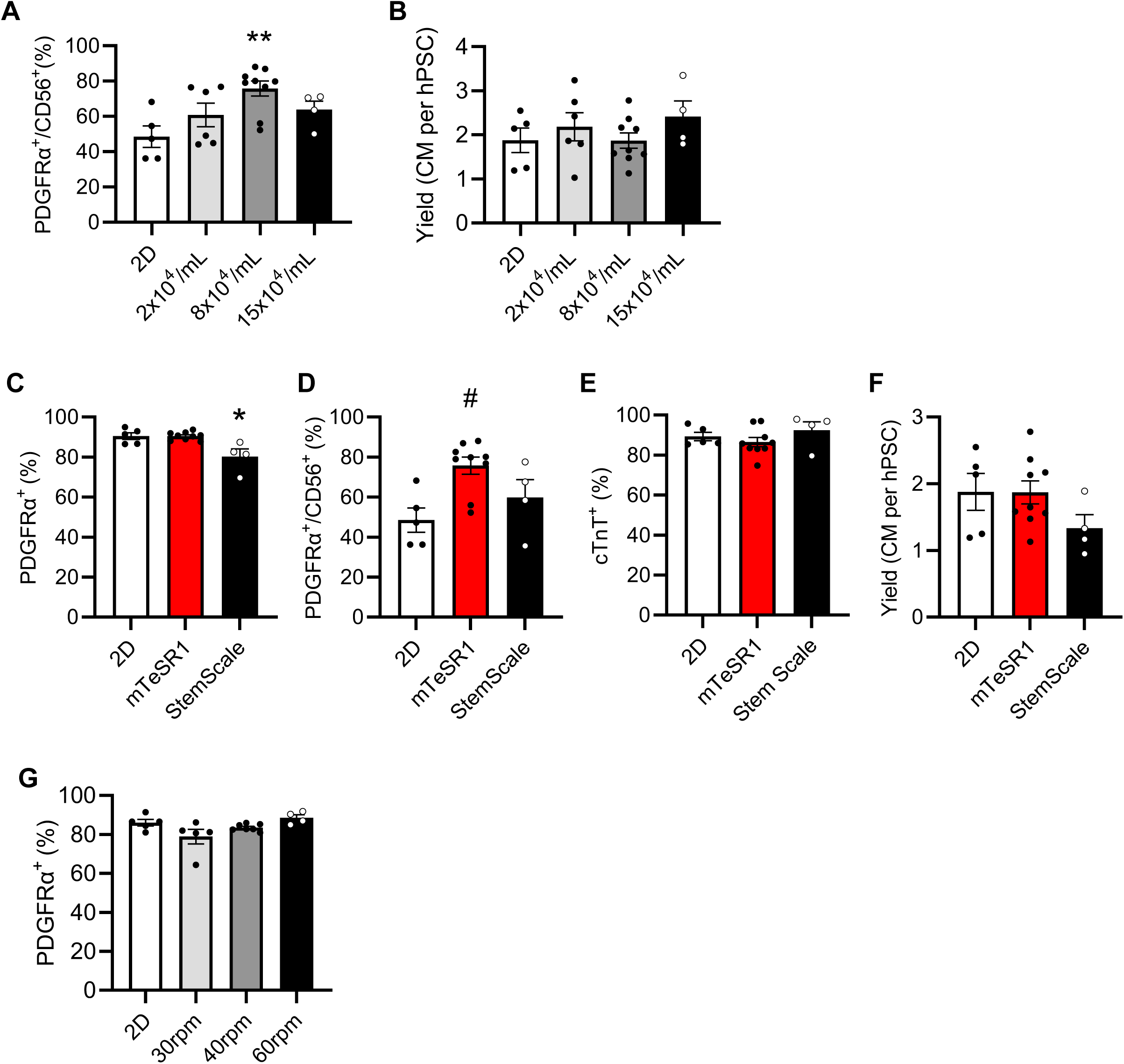
hPSC-CM differentiation optimization. A) Quantification of PDGFRα^+^/CD56^+^ cells on day 3 of differentiation using hPSCs expanded in the VWBR at different seeding densities. B) hPSC-CM yield on day 20 of differentiation. **p<0.05 8x10^4^ cells/mL vs. 2D. C) hPSCs were expanded in the VWBR in StemScale and cardiomyocyte differentiation potential compared to 2D and 3D cells expanded in mTeSR1. Quantification of C) PDGFRα^+^ D) PDGFRα^+^/CD56^+^ cell frequency on day 3 of differentiation. Quantification of E) cardiomyocyte purity (%cTnT^+^) and F) yield on day 20 of differentiation. *p<0.05 3D StemScale vs. all other groups, and ^#^p<0.05 2D mTeSR1 vs. 3D mTeSR1 by one-way ANOVA followed by a Tukey post hoc. G) PDGFRα^+^ frequency on day 3 during differentiation in the 100mL VWBR using hPSCs previously expanded at different agitation rates.

**Supplementary Figure 6.**
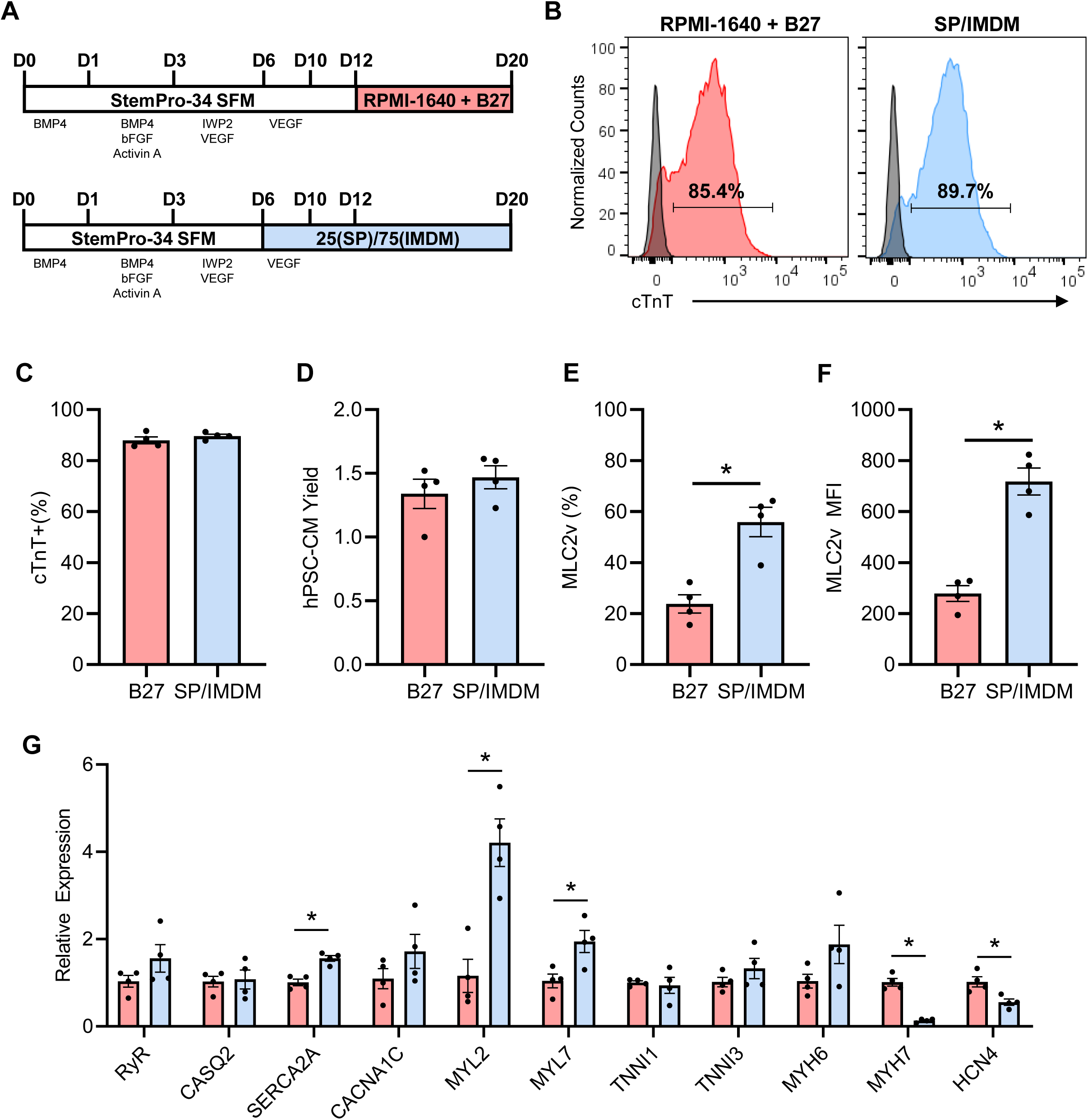
Comparison of RPMI-B27 and SP25 medium for hPSC-CM maintenance. A) Schematic of media used for testing hPSC-CM maintenance. In brief, hPSCs were initially differentiated to hPSC-CMs in 100% StemPro, after which cells were switched to SP25 (25%/75%, v/v; SP/IMDM) on day 6 or maintained on 100% StemPro. On day 12, cells were switched to RPMI-B27 (media condition 1) or maintained on SP25 (media condition 2) until day 20. B) Histogram plots and C) quantification of hPSC-CM purity. D) hPSC-CM yield at day 20 of differentiation. E) Quantification of cTnT^+^/MLC2v^+^ cell frequency and F) MLC2v mean fluorescence intensity (MFI). G) Gene expression in day 20 hPSC-CMs produced using the two different protocols. *p<0.05 by two-tailed Student’s t-test.

**Supplementary Figure 7.**
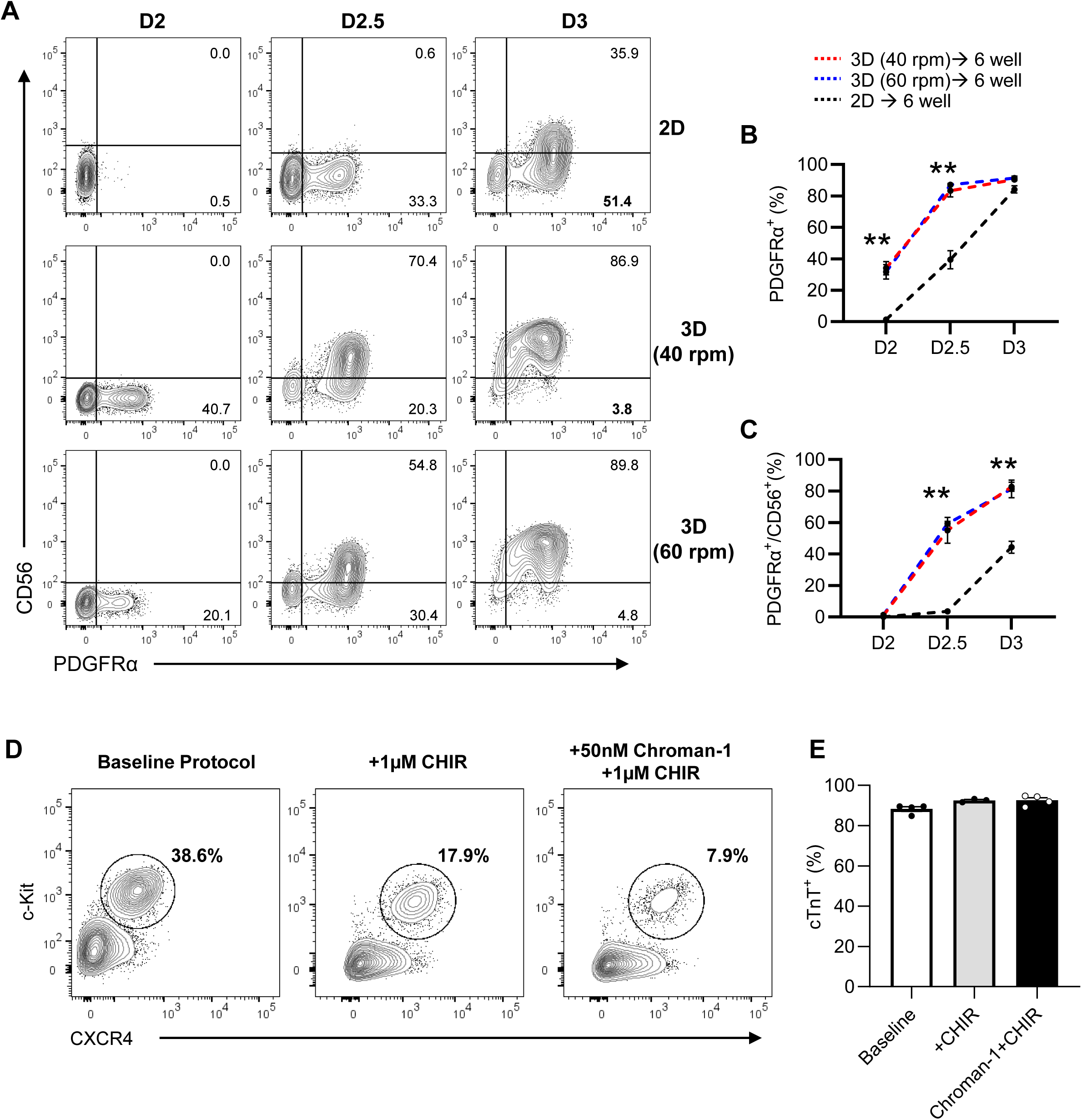
Optimization of hPSC-CM differentiation from VWBR expanded hPSCs. Cells were expanded in either 2D or in VWBRs at 40 rpm or 60 rpm and subsequently differentiated in 6 well plates. Mesoderm induction kinetics were assessed on days 2, 2.5 and 3 of the differentiation. A) Representative plots and quantification of B) PDGFRα^+^ cells and C) PDGFRα^+^/CD56^+^ cells on days 2, 2.5 and 3 of differentiation. D) Flow cytometry plots of c-Kit^+^/CXCR4^+^ populations in the VWBR on day 4 of the differentiation. Plots are shown for the different protocols tested in Figure 4 E-H. E) hPSC-CM purity of cells harvested under the different protocols assessed in Figure 4E-H.

**Supplementary Figure 8.**
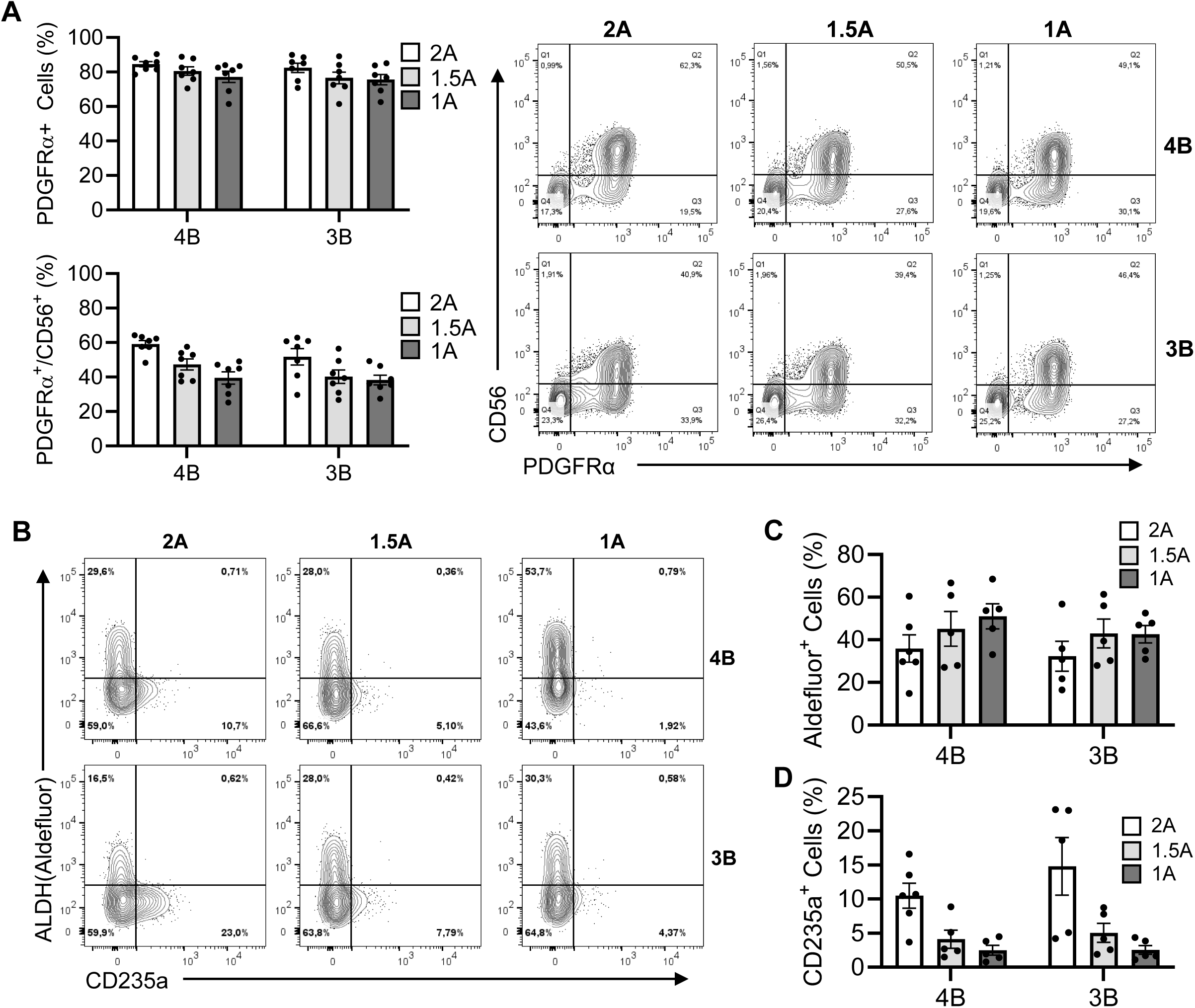
Titration of BMP4 and Activin A levels for optimization of ALDH and CD235a expression in day 4 mesoderm populations. Cells were expanded in the VWBR and differentiated in 6 well plates to determine the optimal cytokine concentrations for second heart field progenitors based on ALDH activity and CD235a expression. A) Quantification of mesoderm commitment (PDGFRα^+^, PDGFRα^+^/CD56^+^) obtained with different BMP4 and Activin A concentrations and representative plots. B) Representative plots showing aldehyde dehydrogenase (ALDH) activity (Aldefluor) and CD235a^+^ expression on day 4 of differentiation and quantification of C) Aldefluor^+^/CD235a^-^ cells and D) CD235^+^/Aldefluor^-^ cells.

**Supplementary Figure 9.**
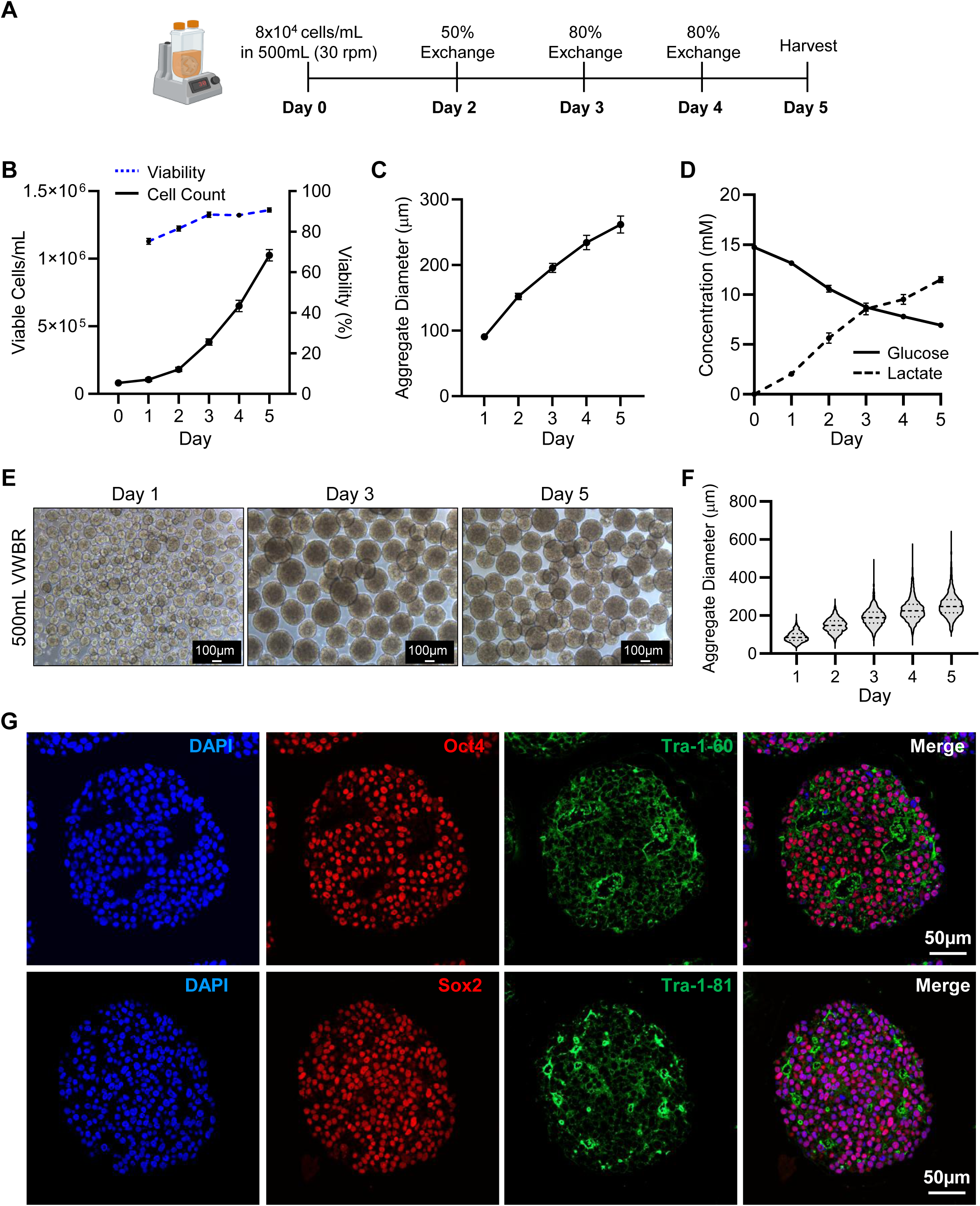
hPSC expansion in 500 mL VWBRs. A) Schematic illustrating the protocol used to expand hPSCs in the 500 mL VWBR. 500 mL VWBRs were seeded with 8x10^4^ cells/mL and cultured for 5 days at 30 rpm using the medium exchange schedule shown in A). Resultant parameters include B) viable cells/mL and cell viability, C) mean aggregate diameter, and D) medium glucose and lactate concentrations over the 5-day hPSC expansion period. E) Representative images of undifferentiated hPSC aggregates during expansion in the 500 mL VWBR. F) Aggregate diameter distribution over the hPSC expansion protocol. G) Representative immunofluorescence images of pluripotency markers in aggregates harvested from 500 mL VWBRs on day 5 of the expansion protocol.

**Supplementary Figure 10.**
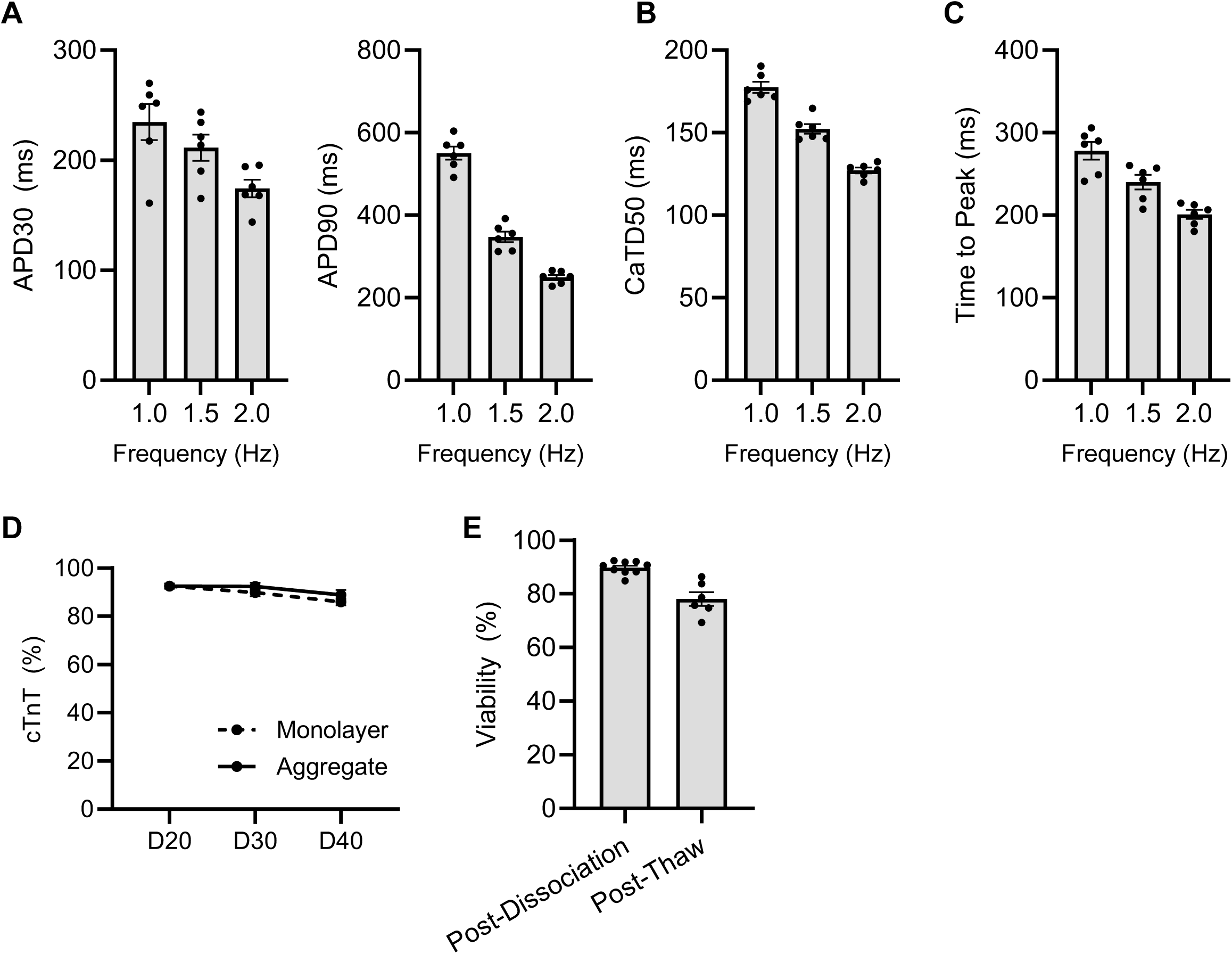
Phenotyping of 500 mL VWBR produced hPSC-CMs. A) Action potential duration (APD) 30 and APD 90 in hPSC-monolayers electrically paced at 1, 1.5, and 2Hz. B) Calcium transient duration 50 (CaTD 50) and C) time to peak in hPSC-CM monolayers paced 1, 1.5, and 2Hz. D) hPSC-CM purity (% cTnT^+^) at day 20, 30 and 40 in hPSC-CM aggregates and hPSC-CM monolayers from 500 mL VWBR produced hPSC-CMs. E) Viability of 500 mL VWBR produced hPSC-CMs following collagenase dissociation on day 20 and following cryo-recovery prior to transplantation.

**Supplementary Table 1.** Primary Antibodies.

| Antibody | Clone | Concentration | Supplier |
| --- | --- | --- | --- |
| Oct4 | C30A3 | 1:200 | Cell Signalling Technology |
| Sox2 | D6D9 | 1:400 | Cell Signalling Technology |
| Tra-1-60 | Tra-1-60(S) | 1:200 | Cell Signalling Technology |
| Tra-1-81 | Tra-1-81 | 1:100 | Cell Signalling Technology |
| cTnT | 13-11 | 1:100 | ThermoFisher Scientific |
| Oct4-APC | 3A2A20 | 1:200 | BioLegend |
| Sox2-PacificBlue | 14A6A34 | 1:20 | BioLegend |
| PDGFRα (CD140a)-PE | αR1 | 1:10 | BD Biosciences |
| NCAM (CD56)-APC | B159 | 1:100 | BD Biosciences |
| c-kit (CD117)-FITC | 104D2 | 1:100 | BD Biosciences |
| CXCR4-BV421 | 12G5 | 1:100 | Biolegend |
| CD235a-APC | GA-R2 (HIR2) | 1:100 | BD Biosciences |
| cTnT-APC | REA400 | 1:100 | Miltenyi Biotec |
| MLC2v-PE | REA401 | 1:100 | Miltenyi Biotec |
| MLC2a-VioBlue | REA398 | 1:50 | Miltenyi Biotec |
| α-actinin | EP2529Y | 1:200 | Abcam |
| Ku80 | C48E7 | 1:400 | Cell Signalling |

**Supplementary Table 2.**
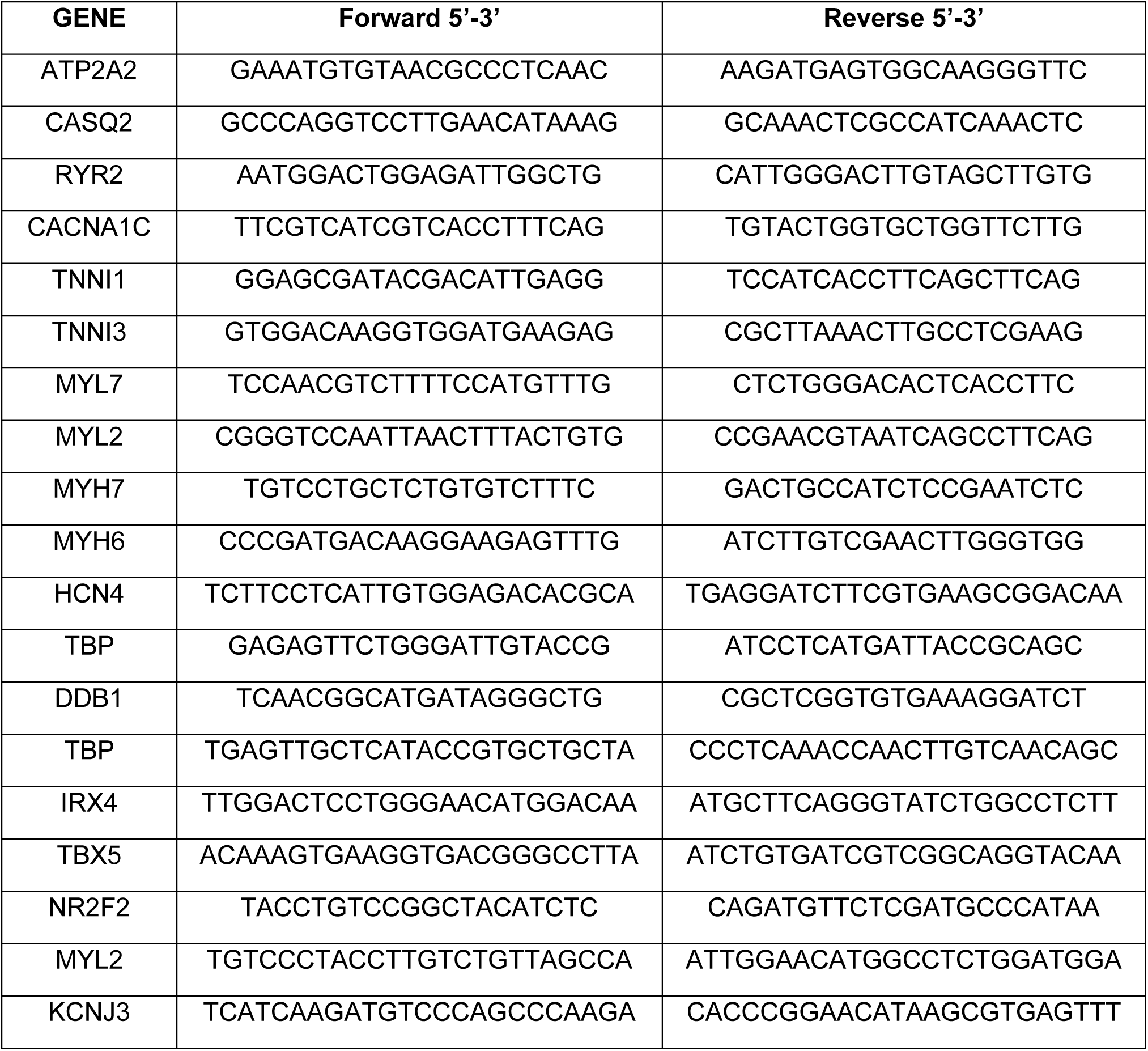
Primer List.

